# Impact and characterization of serial structural variations across humans and great apes

**DOI:** 10.1101/2023.03.09.531868

**Authors:** Wolfram Höps, Tobias Rausch, Peter Ebert, Human Genome Structural Variation Consortium (HGSVC), Jan O. Korbel, Fritz J. Sedlazeck

## Abstract

Modern sequencing technology enables the detection of complex structural variation (SV) across genomes. However, extensive DNA rearrangements arising through series of mutations, a phenomenon we term serial SV (sSV), remain understudied since their complexity poses a challenge for SV discovery. Here, we present NAHRwhals (https://github.com/WHops/NAHRwhals), a method to infer repeat-mediated series of SVs in long-read genomic assemblies. Applying NAHRwhals to 58 haplotype-resolved human genomes reveals 37 sSV loci of various length and complexity. These sSVs explain otherwise cryptic variation in medically relevant regions such as the *TPSAB1* gene, 8p23.1 and the DiGeorge and Sotos syndrome regions. Comparisons with great ape assemblies indicate that most human sSVs formed recently and involved non-repeat-mediated processes. NAHRwhals reliably discovers and characterizes sSVs at scale and independent of species, uncovering their genomic abundance and revealing broader implications for disease than prior studies suggested.

## Introduction

Continuous advances in single-molecule sequencing technologies drive the discovery of increasingly complex patterns of genetic variation in the human genome, particularly in repetitive regions. These highly rearranged regions and the respective complex alleles contribute to population diversity and impact a wide range of phenotypes ^1–4^. Investigating complex alleles in distinct population ancestries is important for elucidating their pathogenic impact, as well as their recent and past evolution. In spite of continued advances in genome assembly, we are far from understanding the full spectrum of human genetic variation particularly in repeat-rich genomic regions, leaving much of their impact on evolution, diversity, and human diseases in the dark ^5–8^.

Structural variation (SVs) in the genomes of humans and other animals are increasingly being characterized at base pair resolution, driven by advances in long read and homolog-preserving genomic technologies ^9–11^. Recent long-read studies have revealed growing numbers of rearrangements of a complexity that escape analysis using conventional short-read sequencing. These rearrangement patterns are caused either by (i) complex genomic rearrangement processes ^12,13^, or (ii) by serial rearrangement events that accumulate at a locus gradually or rapidly due to its proneness to undergo SV ^12,14^. The latter are thought to be particularly predominant in genomic regions harboring segmental duplications (SDs), which facilitate *de novo* rearrangements via non-allelic homologous recombination (NAHR) ^12^. This spatial preference makes their discovery and interpretation especially challenging – with short read based mappers struggling in SD-rich regions – while raising the notion that such SVs may be especially relevant among the fraction of human variation that remains to be discovered ^15^. In light of their ‘serial’ nature, we term the latter *serial structural variants* (*sSV*) throughout this manuscript.

While the frequency of sSVs in healthy and diseased individuals is poorly explored, these regions are highly relevant for population and medical genetics – since they demarcate regions with high diversity in haplotype structure, and regions prone to undergo *de novo* rearrangements including pathogenic copy-number variation (CNV) ^13,16–18^. We recently reported several isolated sSV-like events^14^, which included rearrangements that likely facilitate - or protect against - disease-causing copy number variations in the human genomic loci 3q29, 15q13.3 and 7q11.23. In addition to this, a range of further studies reported medically relevant SD-associated CNVs that could be interpreted as sSVs.

Sanchis-Juan and colleagues proposed likely sequences of mutational events causative for the Coffin-Siris syndrome, cone-rod dystrophy, intellectual disability and seizure and neonatal hypoxic-ischaemic encephalopathy in patients ^17^. Similarly, sSV events potentially causative for early-onset neuropsychiatric disorders ^19^ and Angelman syndrome ^20^ have been identified. From a population genetics viewpoint, the *TCAF1/2* locus displays substantial human-specific sSV-like copy-number variation associated with positive selection and an implicated role in adaptation ^21^ and the *TBC1D3* gene family, similarly, displays remarkable human diversity attributable to sSVs implicated in the expansion of the human prefrontal cortex ^22^. Recently, the first genome-graph released by the Human Pangenome Reference Consortium (HPRC) has allowed new insights into highly variable complex regions, such as the *RHD, HLA-A, C4, CYP2D6* and *LPA*-containing loci, many of which harbor interspersed SDs and are thus likely candidates for sSV activity ^23^. Indeed, for example, SD-associated variation in the LPA locus has been reported to directly impact the risk for cardiovascular diseases ^24^. Yet, despite their undisputed relevance, these regions remain poorly resolved and consequently understudied, owing to challenges in the discovery and interpretation of genetic variation in these regions.

The identification of sSVs using long-read sequencing ^11,25^ or long-read based genome assemblies ^9,26,27^ is conceptually separate from ‘classical’ SV calling^28^, as even a perfect description of sequence alterations (e.g., ‘Del-Inv-Del’) does not necessarily capture the series of underlying simple SVs (such as an inversion followed by a deletion, denoted ‘Inv + Del’ in Figure 1A). Even when this ‘mechanistic’ viewpoint is ignored, only few methods for resolving complex patterns of SV have been developed, and these come with remaining limitations ^11,29^. These limitations include the needs for specialized algorithms designed to capture complex multi-breakpoint SVs thought to be formed through a single mutational event^11^ dependent on the region of the genome these tools are applied to. De novo genome assemblies, which since recently achieve remarkable resolution across nearly the full humangenome^30^, theoretically allow for a more comprehensive study of sSV. However, the identification of sSV from assemblies remains challenging^31^ and to date has not been explicitly addressed through computational methods. Thus, novel methodologies are required to address our current lack of detection and improve our understanding of how sSV contributes to genomic variation.

**Fig 1.**
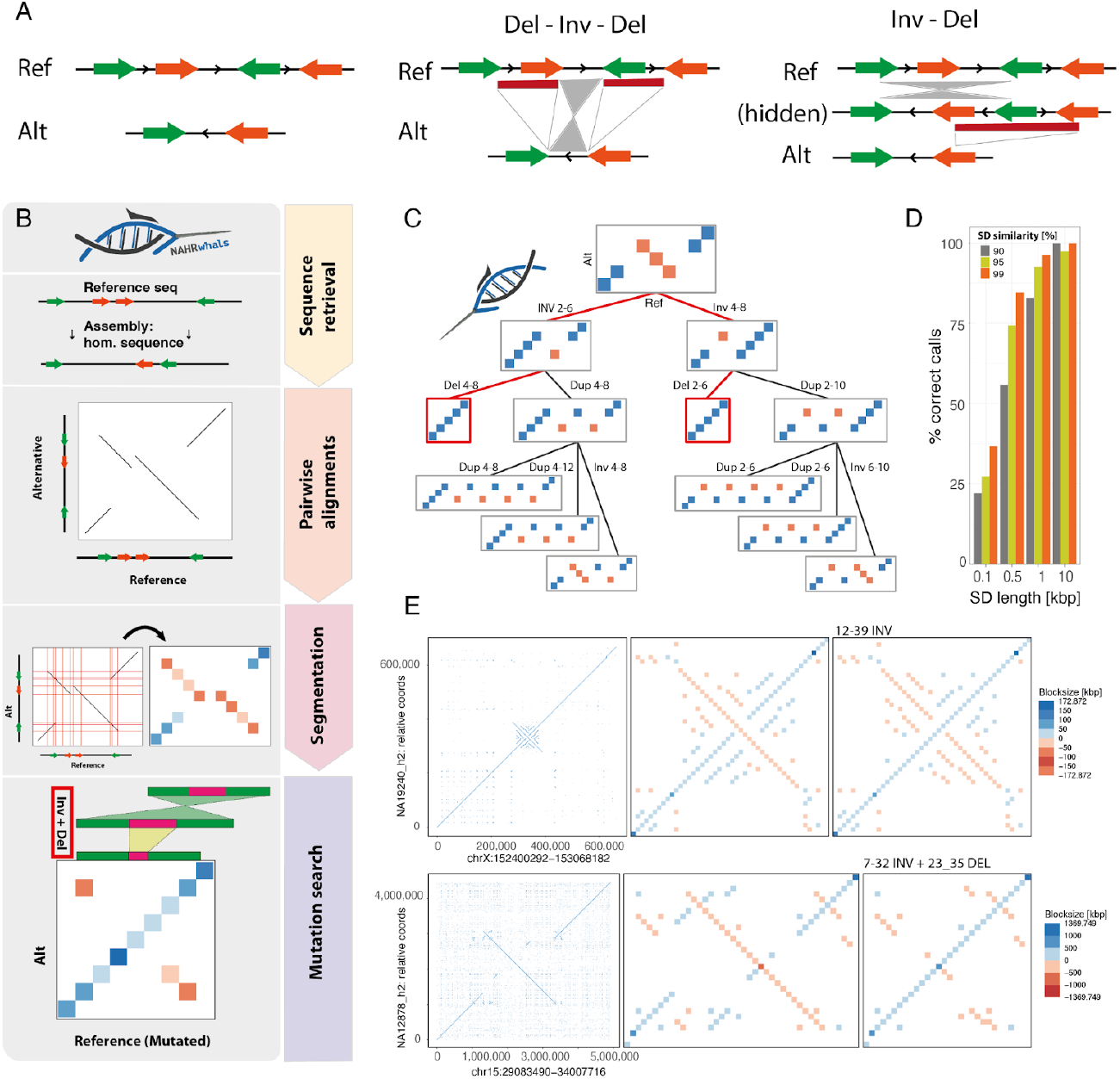
Overview: the NAHRwhals sSV detection method. **A** Schematic representation of a sequence pair illustrating the principle of serial SVs (sSVs). Traditional SV calling is often limited to detangling only simple SV (Del vs. Inv) or to report the entire allele (Del-Inv-Del; middle). Instead, NAHRwhals infers a series of simple SV which can explain a given structural haplotype outcome (Inv-Del; right). **B** Flowchart showing the key steps of the NAHRwhals algorithm. Starting from reference and alternative assemblies and a reference region of interest, the homologous region is first extracted from the assembly. Pairwise alignments between Ref and Alt are created and segmented into a condensed representation. Using this condensed dotplot, an exhaustive search is invoked to explore possible chains of NAHR-mediated rearrangements explaining the structural differences. **C** An example mutation search tree of depth 3 for a simple condensed dotplot. Successful mutation chains are highlighted in red. **D** Results of sSV calling on simulated runs. SD length and similarity correlate with prediction accuracy as longer/more similar sequences are more likely to be retained both in the initial alignment and the dotplot segmentation **E** Two examples of segmentation and mutation calling in real loci. Left: dotplot of pairwise alignments between hg38 (x) and assemblies (y). Middle: Condensed dotplot representation. Right: Condensed dotplot after application of the highest-scoring SV chain.

To directly address this existing gap, we present a computational framework, NAHRwhals (NAHR-directed Workflow for catcHing seriAL Structural Variations), which is based on near-lossless encoding of pairwise genomic alignments, and allowing to infer regions that, based on their local pattern of haplotype structure variation, have undergone series of overlapping SVs (i.e., sSVs). Given a genomic assembly fasta file and genomic reference coordinates of interest, NAHRwhals extracts analogous loci, identifies structural differences and repeat regions, and finally employs an exhaustive search over chains of NAHR-based events that can explain the observed difference in sequence architectures – to generate hypotheses about serial rearrangements resulting in sSVs across the genome. NAHRwhals thereby leverages the sequence resolution of genome assemblies, to enable identification of patterns of complex variation inaccessible to prior methods.

The tool can be readily applied using any genome assembly and a freely exchangeable reference sequence, making it suitable for comparative genomic analysis. Using NAHRwhals and 58 assembled genomes – 12 of which were previously unpublished – we reveal the occurrence of sSV in humans of diverse origin and highlight implications into medically important regions.

## Results

### Automated detection of serial structural variations (sSV) from genome assemblies

To allow systematic characterization of sSVs in haplotype-resolved genome assemblies, we devised the *NAHRwhals* (NAHR-directed Workflow for catcHing seriAL Structural Variations) framework, which identifies regions likely to have undergone sSVs from genome assemblies. NAHRwhals computes a pairwise alignment between a reference and assembly, which is subsequently simplified into representing only the general architecture of analogous, rearranged and repeated segments. Subsequently, an exhaustive mutation search then identifies SV-chains that can transform the reference architecture into a structure equivalent to the mutated version. The main analytical steps carried out by *NAHRwhals* (see also Figure 1B; Methods) are: i) *Sequence retrieval:* Minimap2 ^32^ is used to custom-liftover locus coordinates from a reference (Ref, typically hg38 or CHM13-T2T) to an ‘alternative’ (Alt) assembly (a user-provided de-novo assembled genome), identifying and extracting the most similar homologous region in the assembly. This step can be sped up by using a pre-computed alignment file. ii) *Pairwise alignments:* After isolating a locus on Ref and its homologous counterpart on Alt, minimap2 is invoked to produce an accurate local pairwise alignment. Since minimap2 natively displays difficulties in accurately mapping inversions and repetitive sequences, we devised a custom mapping pipeline in which Alt is split into chunks whose length scales with the total sequence length (default: chunklength [kbp]/upper threshold sequence length [kbp]: 0.1/10, 1/100, 10/5000, 50/Inf) (Methods). The sequence chunks are then aligned to Ref individually, and later re-joined to form long alignments (Methods, Figure S1). iii) *Alignment segmentation:* To prepare segmentation, very short alignments are removed and alignment start-/endpoint coordinates are rounded to the closest multiple of a rounding factor (default: 1-10 kbp depending on sequence length) to eliminate small alignment incongruencies. A custom segmentation algorithm then processes the output of the pairwise mapping by identifying horizontal and vertical overlaps between alignments and defining ‘quadrants’ representing unique sequence motifs of varying length (Figures 1B, S2, Methods). These quadrants can be visualized in the form of a ‘condensed dotplot’ which comprises a faithful representation of the original alignment. iv) *Exhaustive mutation search:* Using the condensed dotplot as a basis, the mutation space is finally explored in a *breadth-first* search approach, with the goal of identifying NAHR-based SV-chains capable of transforming the reference sequence (represented on the x-axis) into a structure equivalent to Alt (Figure 1C). In the space of condensed dotplots, repeat pairs are represented by pairs of colored squares found in the same row, and deletions, duplications and inversions can be simulated very efficiently by deleting, duplicating or inverting columns between such repeat pairs, respectively. A custom evaluation function is used to assess the mutated condensed dotplots and identify successful chains (Methods). Variation is considered as ‘explained’ if the best-scoring mutational chain exceeds a user-defined threshold in sequence congruence established between the Ref and Alt (recommended for human-human comparison: 98%). Due to the computational complexity, in practice we limit the depth-of-search to 3 consecutive mutations. The computational time depends on the sequence length, complexity and depth of search for a given locus and ranges typically between 15-60 seconds, with up to 5 min CPU time in extreme cases.

To assess the performance of NAHRwhals, we created and surveyed 1,200 artificial sequences containing two pairs of segmental duplications each, and simulated serial SVs of depth one to three (e.g. Dup-Inv-Del) (Methods, Figure S3). We additionally randomized the length (0.1 kbp - 10 kbp) and similarity (90% - 99%) of segmental duplications to estimate thresholds above which pairwise alignment are typically reported and represented in the condensed dotplots. We find a positive correlation between genotyping accuracy and the length and similarity of repeats, with near-perfect sSV-detection accuracy if the repeat length exceeded 10 kbp in this simulated setting, while short (100’s bp) or highly dissimilar repeats (<90%) are typically rejected from sSV simulation (Figure 1D). To further confirm the viability of NAHRwhals in a more realistic setting, we applied NAHRwhals to ten previously identified inversion loci of varying complexity and known to be subject to NAHR^14^. In this benchmarking exercise, NAHRwhals obtained accurate condensed representations and expected SV genotypes for all ten loci (Figures 1E, S4). In conclusion, our method shows promising results across simulated and previously characterized NAHR-affected regions along the genome.

### Automated reconstruction of sSVs across 336 loci along the human genome

Having established the general ability of NAHRwhals to infer serial SV through simulations and example test loci, we next performed a broad survey for sSV events in a diverse panel of 56 assembled human haplotypes. These haplotype-resolved assemblies were generated by the Human Genome Structural Variation Consortium (HGSVC) using Pacific Biosciences long read sequencing. These include hitherto unpublished assemblies of 6 human genomes, amounting to 12 assembled haplotypes, generated using 27.7-47.2X coverage Pacific Biosciences long reads (HiFi) that were phased into chromosome-scale haplotypes using Strand-seq ^33^ (Methods). Since these 56 assemblies span human individuals of diverse population ancestry, they allow us to obtain an estimate for the prevalence of sSVs in humans and identify classes of sSV-mediated variation. To this end, we defined a list of potentially sSV-carrying genomic regions by applying a merging strategy in which we integrated: a) all SV regions longer than 10 kbp determined in a previous comprehensive SV survey of 64 human haplotypes ^9^ (n=915); b) sites of polymorphic human inversions ^14^ longer than 10 kbp (n=290); and c) segmental duplications obtained through the UCSC Table Browser ^34^. Using this procedure, we determined 336 non-overlapping loci (median length: 170.8 kbp), which were subsequently scanned for sSV content. In each of these regions, we tested a set of 56 diverse human and four great-ape assembled haplotypes (Methods) for (s)SVs with respect to the CHM13-T2T-v1.1 genome assembly ^30^. Regions were provided in GRCh38-coordinates to NAHRwhals, which converted input coordinates to CHM13-T2T using a custom segment-liftover procedure based on minimap2 (Methods).

By screening across these samples and loci, NAHRwhals inferred 37 loci with at least one instance of overlapping SVs likely to have arisen in a serial manner (Table S1A). Adjacent (i.e. non-overlapping) SVs were not considered as serial SVs. Among the remaining 299 regions, 8 regions did not display a contiguous assembly in any sample and 21 regions could not be explained by NAHRwhals (despite being assembled in at least one sample), indicative of the involvement of non-NAHR mediated mechanisms. All other loci (270) displayed only zero-stage (Ref) or single-stage SVs (Inv, Del, Dup) (Figures 2A, S5). The group of 37 sSVs was subsequently retained for further analysis (Figure 2B, Table S2). Across the 37 sSV regions in 58 human haplotypes, we identified 163 SVs of predicted depth 2 or 3. Notably, 65% of all predicted intermediate states (e.g., ‘Inv’ for an ‘Inv-Del’ haplotype) were indeed observed in another sample, suggesting that most of these complex NAHR loci are the result of accumulation of serial, temporally distinct events. Reflective of the often complex nature of the loci, out of all 2109 sequences (equal to 37 loci in 57 samples), assembly breaks prevented detailed analyses of potential sSVs in almost one third (625/2109 (29.6%)).

**Fig 2.**
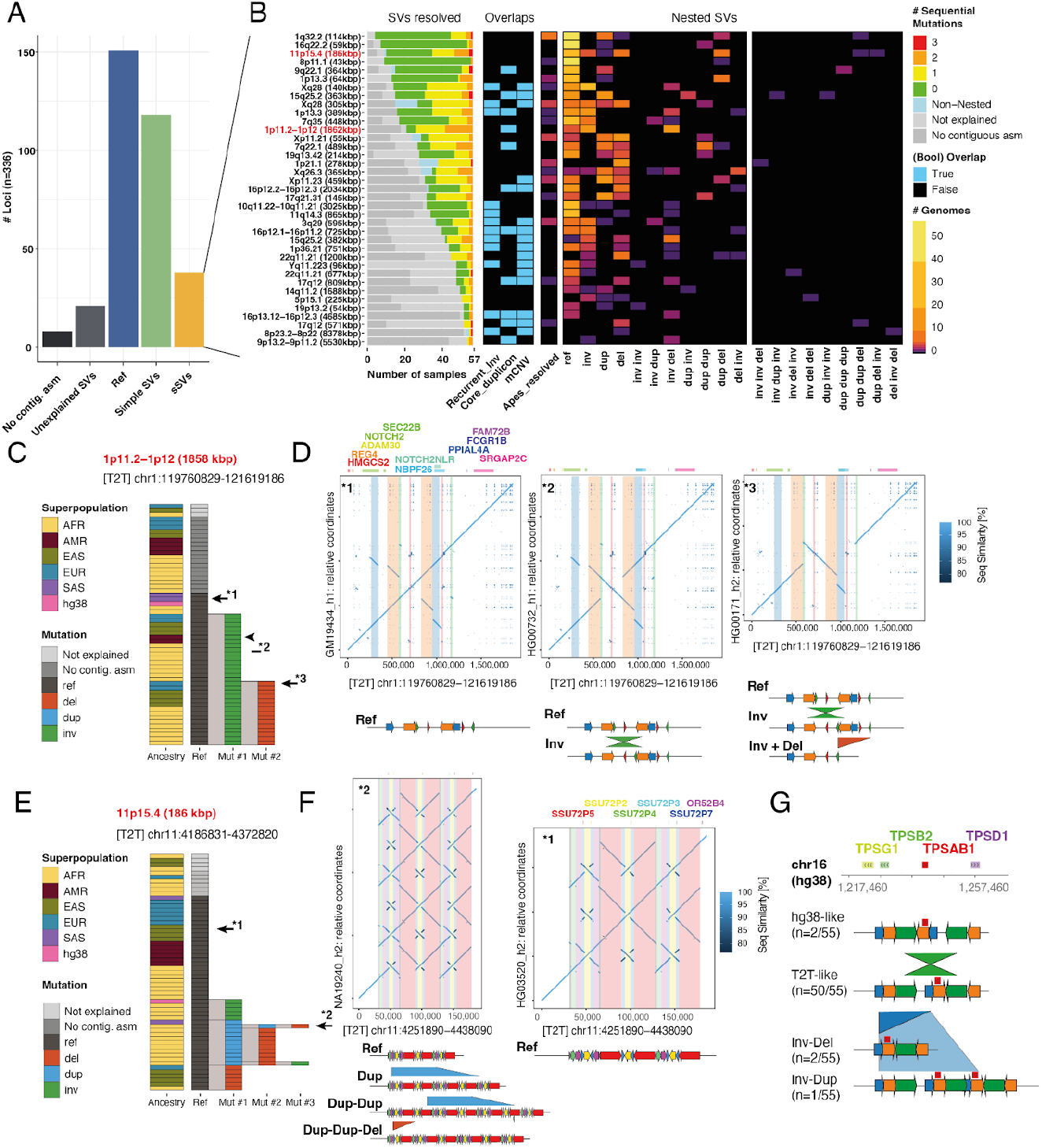
Inversion regions identified as sSVs. **A** Broad classification of the 336 loci initially surveyed with NAHRwhals. Loci were considered as containing sSVs if they displayed at least one overlapping pair of SVs in at least one sample. **B** Overview over the full callset of 37 inversion-containing loci in which sSVs were discovered in at least one sample. The diagram shows the prediction performance in humans and apes (‘SVs resolved’), the presence of recurrent inversions, core duplicon-mapping genes and morbid CNV regions in the genomic region, as well as genotypes for each locus. **C** Three distinct sequence configurations observed in the 1p11.2-1p12 sSV. 15/38 samples harbor a deletion preceded by an inversion compared to CHM13-T2T. **D** Dotplots and SD schematics illustrating examples of all three configurations. **E** A 185 kbp region on 11p15.4 showing complex patterns of nested SVs leading to extreme diversity in the region explicable by NAHR. **F** Dotplot and SD schematics of a highly rearranged (left) and a reference-like (right) alternative configuration. **G** sSVs found in the disease-relevant TPSAB1-containing region. Observed CNVs can be explained as simple SVs on the CHM13-T2T-like configuration, but appear as sSVs with respect to hg38 (Figure S15).

Lastly, 540/2109 sequences (25.6%) were considered as ‘unexplained’ as NAHRwhals did not indicate either a reference state or any NAHR events. We investigated these further by manually curating 35 of these alignments - one for each sSV loci which had at least one unexplained sample (Figure S6). The majority indeed displayed either small-scale (12/35; 34.3%) or large-scale (7/35; 20%) non-NAHR rearrangements. Further 10 regions (28.6%) exceeded the borders of our window or no homologous alignment was found. In six cases (17.1%), NAHRwhals was too conservative in rejecting alignments for exceeding boundaries (n=4) or missed an optimal solution due to prematurely aborting branches of the mutation search tree (n=2).

To better visualize and report the sSVs results across multiple samples, we devised a visualization resembling a directed flowchart, in which each temporally distinct NAHR mutation event is represented by a node (Figure 2C,E). To illustrate the types of sSVs identified with NAHRwhals, we first highlight a ∼1.5 Mbp region on chromosomal region 1p11.2-1p12, which displays several pairs of overlapping SDs in the reference state (carried by CHM13-T2T, hg38 and four other assemblies). A simple inversion between one of the SD pairs was observed in 17 samples, and finally 15 samples were carrying a third configuration which presumably features deletion of the inverted haplotype, corresponding to an ‘Inv + Del’ sSV (Figure 2C, D). Another example of a more complex sSV-rich region was found in chromosomal region 11p15.4. In this case, several distinct haplotype configurations were observed, which could be explained by one, two and three consecutive SVs, respectively (Figure 2E, F). Our callset includes also other regions for which sSV-like patterns have been described previously, such as variation in regions containing *TCAF1/TCAF2* (7q35 (448 kbp))^21^, *POTEM/POTEG* (14q11.2 (1688 kbp)^35^, *TBC1D3* (17q12 (571 kbp)) ^22^, *AMY1A-C* (1p21.1 (278 kbp))^10^ and others (see Table S1A for a full list) – all of which underwent dynamic SD-associated rearrangements in human and great ape evolution ^21^. sSV plots and dotplot visualization for each of the 37 sSVs are in the supplementary material (Figures S7-S14).

The sequences of the 37 sSV loci were additionally validated by mapping ultra-long nanopore reads (generated by the Human Genome Structural Variation Consortium, and available for 11/26 samples) directly to the respective assemblies and applying Sniffles2 ^36^ to scan for homozygous SVs, which would be indicative of likely assembly errors (Methods). Across 796 thusly tested sequences, we find no instances of homozygously rearranged or inverted regions, suggesting absence of major misarranged assembly regions. If we employ a length cutoff of 10 kbp, 45 sequences (5.7%) display homozygous indels which may point at collapsed or duplicated regions. Among screened haplotype configurations which were predicted by NAHRwhals only once, the ratio of indel-containing sequences was only marginally higher (17/19 sequences (10.5%)), suggesting again that most reported calls are biologically meaningful.

We also re-analysed all 119 ‘simple SV’ calls, this time using hg38 as a reference to account for the possibility that certain loci may appear as sSVs only in the context of a different reference allele. This analysis yielded four additional hg38-specific sSVs (Table S1B), three of which displayed ‘Dup-Dup’ alleles indicative of a shorter (or collapsed) allele represented in hg38. In the last identified region, a 49 kbp region on 16p13.3, hg38 represents a minor inverted allele which is predicted to undergo ‘Inv-Del’ (n=2) and ‘Inv-Dup’ (n=1) in three samples, while the same alleles appear as simple SVs when compared to CHM13-T2T (Figure 2G). Notably, the region contains the *TPSAB1/TPSAB2* genes (Figure S15), CNVs of which have been associated with Alpha Tryptasemia, a non-lethal hereditary disease affecting 4-6% of the population ^37^. To our knowledge, the mechanistic background of these CNVs has not been clarified previously. Lastly, we note that, when mapped back to hg38, 7/37 (18.9%) sSV loci display long (> 1 kbp) stretches of hard-masked bases, which prohibit faithful SV reconstruction, highlighting the importance of using a contiguous reference for studying sSV loci. Therefore, our screening reveals a high prevalence of repeat-rich regions for sSV formation and further indicates that complex variation in many dynamic human loci can be readily explained in the framework of sSVs.

### sSV occurrence and SV complexity across Hominidae

Having identified abundant sSV loci in humans, we next set out to examine to which degree sSVs in these regions are human-specific, as a prerequisite to gaining an understanding as to how these regions may have emerged evolutionarily. Repeat-associated variation is known to have contributed substantially to the evolution of modern humans ^38^, and hundreds of genomic regions display SD-mediated inversions between non-human primates and humans ^39–41^, frequently accompanied by secondary copy-number variations (CNVs) near their breakpoints ^41^. We considered that a fraction of our human sSV loci, too, may have undergone substantial restructuring during great ape evolution, which may be explicable through sSVs. To test this, we turned our attention to sSVs in the four great ape genome assemblies^22^ included in our dataset (Methods). To account for the overall higher sequence divergence, we chose a lower threshold parameter of 95% for considering sequence reconstruction successful. When reviewing the set of 37 CHM13-T2T-based human sSV loci, we find that in 16/37 loci (43.2%), NAHRwhals could determine rearrangements translating from the human locus configuration to that of at least one great ape variant (Figure 3A).

**Fig 3.**
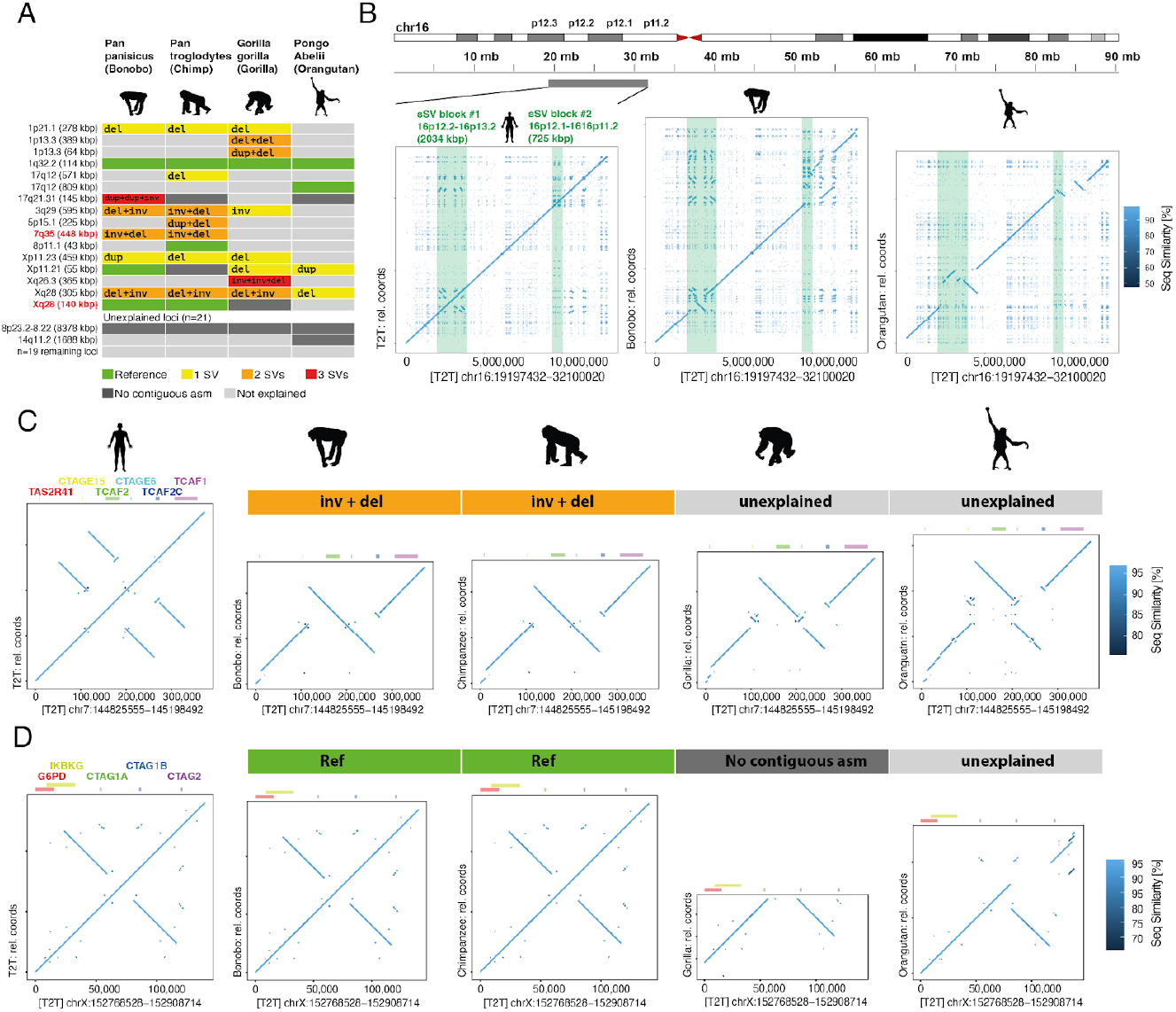
Human sSV loci in great ape genome assemblies. **A** Tabular view of NAHRwhals-based SV genotypes across 37 human sSV sites in four ape species. In 16 loci, the observed variation could be explained for at least one ape haplotype. **B** Dotplot views of a ca. 12 Mbp region on 16p, comparing the assemblies of CHM13-T2T, Bonobo and Orangutan (all single-contig; Chimpanzee and Gorilla: no contiguous asm). Two human sSV loci are highlighted in green. **C** Variation observed in the TCAF1/2 containing region on 7q35. Gorilla and Orangutan likely display a mixture of NAHR and non-NAHR SVs. **D** View of the CTAG1A/CTAG1B containing locus. Compared to Orangutan the CHM13-T2T-, Bonobo and Chimpanzee sequences harbor a duplication, which can not be explained by NAHR.

Out of 37 loci, only four suffered from a lack of locus-spanning contigs in any of the ape assemblies, suggesting that the high ratio of unexplained variants (21/37 regions) does not stem primarily from a lack of assembled sequence quality, but may instead be attributed to non-NAHR-associated SVs or missed calls by NAHRwhals. To decide which of these is the case, we examined dotplot visualizations of these regions (Figures S16-S18). Indeed, the unexplained loci consistently displayed advanced levels of rearrangements, frequently featuring large insertions, deletions and translocations, unattributable to NAHR, which are likely the result of other formation mechanisms, including duplicative transposition events^42^ not modeled by our framework. We illustrate the scope of complexity of sSV loci in great apes along a 12 Mbp region on the p-arm of chr16 (Figure 3B), which harbors two human sSV loci, neither of which could be explained in any great ape assembly. A dotplot visualization of these regions in the two contiguous ape assemblies (bonobo and orangutan) reveals that both sSV loci are part of larger, highly complex rearrangements that exceed the scale of the human sSV both in size and complexity.

In line with greater evolutionary distances involved, we notice a roughly 2-fold enriched fraction of 2- and 3-step vs 1-step SVs in apes compared to humans (simpleSVs/multi-stepSVs/fraction: humans: 302/163/1.85, apes: 11/12/0.92). We again highlight two examples of sSV loci here. First, the aforementioned *TCAF1/2*-containing region on 7q35 displays a set of overlapping SDs in the CHM13-T2T-configuration, which can transition into an “Inv+Del’’ state in Bonobo and Chimpanzee (Figure 3C). The further distant species Gorilla and Orangutan display a somewhat analogous configuration, but also harbor additional insertions that cannot be explained by NAHR alone. We also find instances of more isolated non-NAHR events, such as a duplicated section on Xq28 containing cancer/testis antigen 1 (*CTAG1A/CTAG1B*) genes, which are implicated with a variety of cancers ^43^. In our dataset, the CHM13-T2T assembly, Bonobo and Chimpanzee share the same locus configuration, which is distinct from the non-duplicated region in Orangutan. NAHR was again not sufficient to explain the implicated duplication event. Across these examples and the remaining dataset, our results suggest that in the majority of human sSV loci, NAHR alone is insufficient to explain inter-species variation, where consequently also other mutational mechanisms are likely at play.

### sSV regions co-locate with disease-causing CNVs, core duplicon genes and recurrent inversions

NAHR-mediated recurrent inversions co-cluster with disease-causing CNVs in some of the most dynamically evolving regions of the human genome ^14^. To anticipate whether our sSVs may be able to explain some of the variation in these regions, we initially tested sSV regions for spatial co-location with recurrent inversions and disease-associated CNVs (morbid CNVs) (Table S3). In line with the tendency of (NAHR-promoting) SDs to flank morbid CNV sites ^44–46^, 16/37 sSVs (43.2%) overlapped with a morbid CNV region or its close surrounding (plus/minus 25% of the CNV length), compared to 51/299 (17.1%) of non-sSV loci, corresponding to a 3.7-fold enrichment of sSVs among morbid CNV-overlapping loci (p = 4.9×10^−4^, one-sided Fisher’s Exact Test). To account for the possibility that this effect may be driven by locus size (i.e., larger loci may be more likely to exhibit sSVs while also being more likely to overlap morbid CNVs), we repeated the experiment, this time measuring if the midpoint of a locus falls into a morbid CNV. This did not alter the overall trend, with 15/37 midpoints of sSV-midpoints and 47/299 non-sSV-midpoints overlapping morbid CNV, respectively (odds ratio 3.63, p = 7.1×10^−4^, one-sided Fisher’s Exact Test). Furthermore, 11/37 (29.7%) of sSV loci overlapped at least one member of a gene family mapping to core duplicons such as *GOLGA* and *NPIP* (methods, Table S3), corresponding to a 6.6-fold enrichment compared to non-sSV loci where this was the case for 18/299 (6.0%) regions (p = 5.6×10^−5^, one-sided Fisher’s exact test). This enrichment is in line with the role of core duplicons which are implicated in the expansion of segmental duplication and further repeat-driven mutational processes ^47^. Lastly, sSVs were 6.25-fold more likely to overlap with recurrent inversions than non-sSV loci (12/37 (32.4%) vs 21/299 (7.0%) overlaps; p=3.78×10^−5^, one-sided Fisher’s exact test), supporting the notion that recurrent inversions are disproportionately associated with complex variation ^14^. When again only sSV-midpoints were considered, the enrichment dropped to 3.66-fold (6/37 vs 15/299 overlaps, p=0.019, one-sided Fisher’s Exact Test). Consequently, our screening suggests that, among the 336 initially included loci, sSVs are strongly (3.66 to 6.6 - fold) enriched in regions containing recurrent inversions, morbid CNVs and expanding SDs.

We proceeded to explore the molecular underpinnings of the association between inversions, morbid CNVs and sSV loci in our data. Indeed, by taking into account a larger window around morbid CNV-associated sSV loci, we were able to identify likely associations again using NAHRwhals as an sSV caller. The first case discovered this way lies in chromosomal region 22q11, which can harbor local duplications and deletions associated with the DiGeorge Syndrome. The region contains a network of segmental duplications which are highly variable across humans and thought to represent mediators of the 22q11.2 deletion syndrome ^48^ (Figure 4A). One sSV locus maps to a 1,200 kbp block of SDs flanking the region. The sSV features two pairs of overlapping pairs of inversely oriented SDs, out of which three divergent haplotypes emerge (Figure 4B), two of which lead to expansion or contraction of the SD content, possibly increasing or decreasing the risk for subsequent formation of the larger disease-causing CNV. Our validation of this region with ONT reads (Methods) was not entirely conclusive, suggesting that further variants may be present in this region (Figure S19A). However, the presence of three instances of the predicted intermediate (inverted) state in other samples (Figure 4B) does support the emergence of ‘Inv-Del’ and ‘Inv-Dup’ haplotypes.

**Fig 4.**
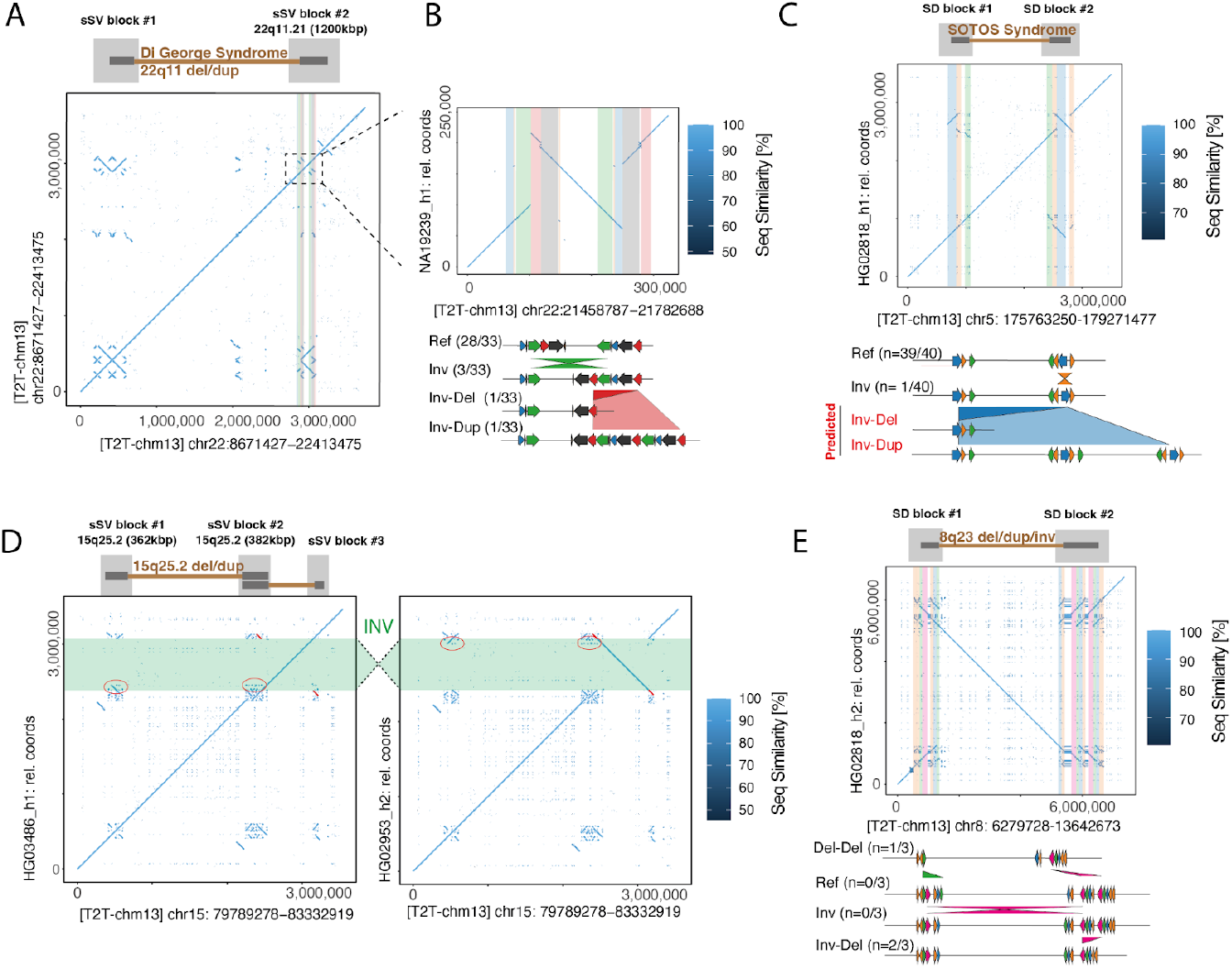
sSVs in disease-relevant regions. **A** Dotplot view of the 22q11 Del / Di George Syndrome region, with two SD-rich blocks of sSV activity highlighted above. **B** Nested SVs lead to duplication or deletion of segmental duplications bordering the 22q11 Del/Dup Di George Syndrome region; potentially affecting the risk of subsequent CNV formation. **C** Inversion of one breakpoint of the SOTOS deletion creates a long pair of directly oriented SVs likely predisposing to subsequent CNV formation. **D** Two sSVs map to two ends of the 15q25.2 deletion region, making both breakpoints susceptible to individual rearrangements. An inversion in one sample leads to transfer of SD sequence to a third sSV block. **E** Variation observed in the highly recurrent 8p23.1 Inv/Del/Dup region. Among the three resolved instances of this locus, NAHR-based variation was observed in both flanking SD blocks.

Primed by this initial example, we set up a more systematic screening to identify sSV events potentially associated with morbid CNV formation. To this end, we investigated a set of 113 disease-associated CNV locations ^44,49^, which we again subjected to a merging procedure to unify redundant calls and include surrounding SDs (Methods), leading to 48 non-redundant morbid CNV locations of interest which were tested with NAHRwhals across all 58 haplotype assemblies (Table S4). Using this procedure, 35/48 loci displayed only reference states, non-contiguous assemblies or unexplained variance. Nine loci displayed single SVs, and another four displayed sSVs of depth 2 (Figure S20, Table S4). Focussing initially on SVs of depth 1, we infer a novel association between an inversion and formation of a ∼2 Mbp CNV associated with the SOTOS syndrome which is likely caused by haploinsufficiency of the NSD1 gene contained in this region ^50^. Repeats at the flanks of this region have been noted before as substrates for NAHR ^50 51^. Our inspection reveals an additional novel 258 kbp inversion of parts of the distal segmental duplication, present in 1/40 resolved haplotypes (2.5%) (Figure 4C) and confirmed by orthogonal ONT-based validation. This inversion flips a substantial SD-containing segment (denoted as DLCR-A in previous literature ^51^) into direct orientation with its proximal counterpart, putting the inverted configuration at risk for subsequent morbid CNV formation.

Turning our attention to the four loci displaying sSV activity (regions 16p12.2-16p11.2 (12.4 Mbp), 15q25.3-15q25.2 (3.7 Mbp), 8p23.1-8p22 (6.9 Mbp), 3q29 (2.6 Mbp); Table S4), we identified further modes of sSV-associated mutation. Since we described variation in the 3q29 region before, we here did not further focus on this region ^14^. In the case of 15q25.3-15q25.2, we observed a ca. 700 kbp inversion which contains further SD fragments and effectively acts as a transporter of those SDs between two SD blocks (red circles, Figure 4D). We also observed sSV activity in the 8p23.1 Inv/Del/Dup region in three resolved samples. Variation in this case is found in both flanking SD blocks (del-del) as well as between the blocks (Inv-Del) (Figure 4E). All assembly configurations described here could be validated with ONT reads (Figure S-19B-D). Extrapolating from these examples, we propose a hierarchical model of sSVs, in which individual ‘blocks’ display variations within themselves, but, additionally, sequence is prone to be exchanged between blocks with inversions as the transport media.

## Discussion

We present NAHRwhals, a long-read assembly-based workflow for identifying and deconvoluting regions of overlapping NAHR-mediated rearrangements. NAHRwhals works by determining local alignments of a region of interest to a reference, and subsequent segmentation of the alignment to summarize syntenic blocks via ‘compressed dotplots’. This massive reduction in data complexity allows us to apply an exhaustive search strategy in which we effectively test all possible sequences of NAHR events up to depth 3 (or higher; at the cost of computation time). Thus, our method closes an essential gap in the current methodology to enable the automatic detection and study of genome complexity caused by overlapping repeat-mediated rearrangements. NAHRwhals can be easily applied to different mammalian species, as we highlighted over the great ape study, to further deepen our understanding of the impact on these complex regions.

We showcase the advances made by NAHRwhals by screening across 46 previously published^52^ and 12 novel haplotype-resolved human genomic assemblies, revealing NAHR-mediated complexities in 37 loci and suggesting that these patterns are far more common in healthy individuals than current genomic studies have suggested ^9^. Among all inferred serial SV (sSV) configurations, roughly two-thirds of predicted intermediate states could indeed be observed in other samples, supporting the notion that the majority of such complexities have formed via serial accumulation of overlapping simple SVs, rather than by individual complex events. It should be noted that NAHRwhals does not incorporate information about population-based haplotype structures in its analysis. As a result, actual locus histories may be more complex than what is predicted by the tool. In evolutionary studies, we recommend validating the predicted series of SVs using complementary methods to gain a more complete understanding of the locus history. Moreover, individual non-NAHR based complex rearrangement events can lead to highly complex patterns of variation, too, ^53^ and these may in some instances be hard to distinguish from sSVs, especially in cases where intermediate haplotype structures are missing. Yet, the presence of flanking segmental duplications at the boundaries of the SV events studied here support a major role of NAHR. In theory, it is also perceivable that serial NAHR events can occur instantaneously as part of the same mutation event, rather than gradually, however such observations have not yet been documented to our knowledge.

When extending our search to greater evolutionary distances, we note that NAHR alone is rarely sufficient to explain variation in sSV loci across four great ape assemblies. Assembly gaps did not contribute significantly to these unexplained variants, supporting the notion that over evolutionary timespans (mya), human sSV loci have seen large-scale rearrangement processes mediated by mechanisms distinct from NAHR. We note that in the future, greater numbers of high-quality ape assemblies will help paint a finer picture, especially concerning the phenomena of incomplete lineage sorting and recurrence^40^.

Human NAHR-mediated complexities are strongly (3 to 6-fold) enriched in regions containing recurrent inversions and disease associated CNVs and can explain variation in medically relevant genes such as *TPSAB1*. sSVs in particular show a dynamic interplay with sequences known to be at risk for CNVs. Such interactions are not unexpected given that morbid CNV regions are frequently flanked by, and nested in, complex repeat patterns which mediate their formation ^14,45,46,54^. We identify at least two modes of interaction. Firstly, the individual flanks of such regions are prone to harboring sSVs, amplifying - or reducing - the SD content available for formation of the ‘main’ CNV. In other cases, we also observe ‘hierarchical’ events in which large inversions span regions of CNV risk and contribute to exchanging sequence between the flanks, putatively increasing their complexity over time, or creating new hotspots of interspersed SDs which can again diversify over time.

NAHRwhals is sensitive to the choice of reference sequence. We showcased this by comparing GRCh38 and CHM13-T2T, where the former suffered from unresolved sequences. These two references frequently represented alternative versions of alleles, some of which are likely attributable to falsely collapsed or duplicated sequences in GRCh38 ^55^, whereas in other cases they may be attributable to repeated mutation and high variability. As highlighted, the dynamic nature of these sequences is especially important as many segmental duplications have an impact on medically relevant genes or other important phenotypes ^22^ and thus motivate a close analysis also in other organisms. To accomplish this, NAHRwhals is able to be adjusted to searches across species or even outside of humidea.

For obvious reasons, NAHRwhals is also reliant on the correctness of the assembly itself. For this study we have assessed the correctness of most assemblies by realigning ultra long ONT reads to themselves and screening for homozygous SV across them. Such SV would indicate assembly errors such as rearrangements that would impact the results of NAHRwhals. Over the screened assemblies, no mis-arragements were observed, and indels indicative of collapses or duplications were rare, too (∼5% of sequences) (Figure S-ont_cnv_validation).

Another important point to accomplish is the automatic parameter tuning that is happening in NAHRwhals. Firstly, we observe a general robustness of the dotplot encryption. Still, our level of compression scales with overall sequence length, meaning that SV predictions can be sensitive to the size of the region of interest – i.e., SVs that affect only a small portion of the window (<10%) can be missed in some cases. Furthermore, by default we consider sequence variation as ‘explained’ if mutated alignments show more than 98% congruence, again discarding variants much smaller than the sequence window. It is likely that a portion of sSVs adjacent to very long CNV sites may have been missed by our survey in this way. Thus, local rearrangements may appear to be unexplained from a narrow angle, but may be explained when the broader sequence context is taken into account.

Overall, this work demonstrates the significance of complex NAHR-shaped variation and its ubiquitous detection across newly assembled genomes, regardless of species. These regions have been demonstrated to be of critical importance in various phenotypes, including those related to disease.

NAHRwhals allows for the automatic detection and study of these regions across multiple assemblies, as well as the identification of the events that likely lead to the complex patterns that are currently observed. With the rapidly increasing number and quality of novel genome assemblies, we anticipate that NAHRwhals will be instrumental in uncovering the origins of disease-causing variants in patients and advancing our understanding of the evolution of these highly variable regions of the genome.

## Methods

### Pairwise sequence alignments

To obtain accurate pairwise alignments even in highly repetitive genomic regions, a custom pipeline was built around the minimap2 aligner (version 2.18) ^32^ to create pairwise alignments: Before aligning, the *query* sequence is split into chunks of 100 bp (if length(query) < 10 kbp), 1 kbp (if length(query) < 50 kbp), 10 kbp (if length(query) is between 50 kbp and 5 Mbp) or 50 kbp (if length(query) > 5 Mbp). The ‘chunks’ are then aligned to the target sequence separately (using the minimap2 parameters *-x asm20 -P -c -s 0 -M 0*.*2*; see Figure S1), reducing the need for read-splitting, which is known to be error-prone in minimap2. The choice of chunklength represents a tradeoff, as a) too small chunks tend to produce overly interspersed alignments which lead to long computation time and a tendency for shorter alignments, and b) overly large chunks tend to ignore or over-merge short alignments. In practice, alignments have proved relatively robust towards the choice of chunklength (Figure S21), justifying our default choices. In a post-processing step, alignment pairs are concatenated whenever the endpoint of one alignment falls in close proximity to the startpoint of another (base pair distance cutoff: 5% of the chunk length). If multiple alignments ‘compete’ for the same partner (e.g., two alignments ending close to the beginning of another), only the longest ‘competitor’ alignment gets selected for merging.

### Noise-reduction in pairwise alignments for subsequent segmentation

Pairwise alignments are retrieved from minimap2 in .*paf* format, in which each alignment can be interpreted as a two-dimensional vector from start (query-start/target-start) to end (query-end/target-end) coordinates. To prepare subsequent compression steps, alignments are pre-processed in multiple ways: First, alignments are filtered by a minimum length threshold (*l*), removing very short alignments. Second, alignment breakpoint coordinates are rounded in x and y direction to the closest multiple of a rounding parameter (*r*). Finally, alignment vectors are shortened along the x or y axis in case that they do not have a slope of exactly 1 or -1 until they do so. The choice of min_aln_len and rounding_factor directly influence the expected dimensionality and complexity of the condensed dotplot. To maximize information content while limiting the size of condensed dotplots, we set l and r to be stepwise functions of the sequence length:

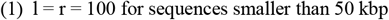

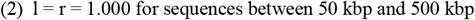

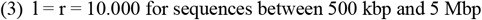

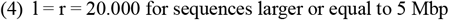

#### Alignment segmentation and dotplot condensation

Following noise-reduction, borders, or *‘gridlines’*, separating unique sequence blocks, are inferred in an iterative way. (Figure S2). In the first iteration, horizontal and vertical gridlines are drawn starting from each start-and endpoint of any alignment. In every subsequent step, overlaps between existing gridlines and alignments are determined, with the points of overlaps serving as a new source for spawning a new *gridline* in perpendicular direction. This process is repeated until no new gridlines are being spawned. In rare cases where the determined grid exceeds predefined maximum dimensions (default: n_rows + n_cols <= 250; n_alignments <= 150), the parameters for minimum alignment length (l) and rounding (r) are doubled until dimension requirements are met. Once the ‘grid’ is established, the length and directionality of each alignment passing a cell are calculated and transferred into a simplified ‘condensed dotplot’ matrix (Figure S2B).

#### Exhaustive mutation search

The massive information reduction obtained from condensing potentially multi-Mb alignments to a matrix of much smaller dimensions allows us to employ an exhaustive search strategy to identify chains of SVs capable of transforming the Ref-configuration into Alt. We focus on exploring NAHR-mediated SVs (deletions, duplications and inversions), and a key observation is that condensed dotplots retain information of repetitive sequences (i.e. rows or columns with >1 colored square). Starting from one condensed dotplot, a recursive *breadth-first* algorithm initially identifies ‘repeat pairs’ in Ref (pairs of colored squares in the same row) and deducts possible NAHR-mediated SVs (Del/Dup between similarly colored squares; Inv between opposites). Subsequently, every SV is being simulated by deleting, duplicating or inverting respective columns of the condensed dotplot, and new higher-depth SVs are inferred and simulated as the search progresses. Mutated matrices are assigned an alignment score using a customized Needleman-Wunsch algorithm ^56^ in which we treat the condensed matrix like a regular pairwise sequence alignment. SV-chains producing a pairwise alignment of sequence identity larger than a threshold (default: 98%) are considered successful.

#### Simulation experiments

We implemented a framework to simulate and mutate SD-containing sequences (available at https://github.com/WHops/NAHRwhals/blob/main/scripts/simulation-dev.R). For each pair of SDs parameters of three different similarity values (90%, 95%, 99%) and four different lengths (100 bp, 500 bp, 1.000 bp, 10.000 bp), we created 50 genomic sequences with 2 pairs of non-overlapping SDs of randomized position and orientation each. Subsequently, we simulated a pool of sequence derivatives, realizing all NAHR-concordant combinations of Inv, Del and Dup up to depth 2. These mutated sequences and their unmutated ‘ancestor’ were given to NAHRwhals for SV calling and results were compared with the known background of sequences.

#### Collection of 336 sSV-regions of interest

In order to maximize the scope of our survey, we based our sSV search on a set of all (n=107,590) SV regions from a previous large-scale SV survey of 64 human haplotypes ^9^, and additionally considered polymorphic inversion calls (n=399), many of which are known to be associated with complex variation ^14^. Given that individual SV calls may sometimes be part of the same sSV block. We devised a specialized strategy to merge individual, spatially nearby SVs into broader ‘SV;Repeat-containing’ regions using the following procedure: (1) filter variants to length >10 kbp to exclude the bulk of non-NAHR events. (2) merge SVs and Inversions with any overlapping segmental duplications (to include mutation-mediating SDs in the loci). (3) Merge any intervals if they have at least 50% overlap. (4) Elongate each region by 25% of its length on either end. (5) Subtract centromeres and ALR repeat regions. (6) Merge regions with <100 kbp distance to each other (7) filter to regions >40 kbp. The resulting regions contain between one and several thousand SV/Inv calls, with region lengths ranging from 40 kbp to 26 Mbp (median: 170.8 kbp). The scripts and all input data are available at https://github.com/WHops/NAHRwhals_rois.

#### Collection of 48 morbid CNV-regions of interest

Loci containing disease-associated CNVs were based on four separate lists from ^44^ (Table 1; 44 regions. Table 2: 14 regions) and ^49^ (Table S2; 19 regions. Table S3: 36 regions), totalling at 113 loci. All regions were transformed from hg18- to hg38 coordinates using UCSC liftover ^57^. Next, we applied the same 7-step merging strategy as used for defining 336 SV-Inv-sSV regions (see methods:Collection of 336 sSV-regions of interest.), resulting in 48 nonredundant regions of interest which were subsequently used for analysis.

#### Human and great ape assembled haplotypes

We based our analysis on 58 human and four great-ape assemblies. The human haplotypes consisted of the GRCh38 and CHM13-T2T assemblies and 56 de-novo assemblies based on PacBio HiFi reads produced by the HGSVC consortium (see Data availability) as previously described (Ebert et al)^9^. Samples considered for assemblies were of diverse ancestry, including individuals from five major superpopulations (African: 16 individuals, Ad Mixed American: 3, European: 4, South Asian: 1, South East Asian: 4), Phased genome assemblies were created in two batches with slightly different procedures:

The first batch (14 / 28 samples, HG00512, HG00513, HG00514, HG00731, HG00732, HG00733, HG02818, HG03125, HG03486, NA12878, NA19238, NA19239, NA19240, NA24385) was produced by an improved version of the PGAS pipeline ^9,58^ (PGASv13). Briefly, an non-haplotype resolved assembly was created with hifiasm v0.15.2 ^59^ after removing potential adaptor contamination in the HiFi reads. This draft assembly was then used as reference in subsequent steps employing Strand-seq data to cluster assembled contigs by chromosomes and to create a phased set of genomic variants during the so-called integrative phasing step of PGAS. The phased set of variants then informed the haplotagging of the HiFi reads, leading to two haplotype-specific read sets per sample. The final phased assemblies were then created by running hifiasm on each haplotype-specific read set. Basic characteristics of the phased assemblies were reported in previous work (see Ebler et al. 2022 ^52^, Supplementary Table 1).

Assemblies of the remaining 14 samples (GM19129, GM19434, HG00171, HG00864, HG02018, HG02282, HG02769, HG02953, HG03452, HG03520, NA12329, NA19036, NA19983, NA20847) were produced by hifiasm v0.16.1-r375 ^59^ using PacBio Hifi data of 27.7 - 47.2X coverage. For 4 samples with parental short read sequencing data ^60^, we used the trio binning assembly mode of hifiasm (GM19129, HG02018, NA12329, NA19983) and for 6 samples with paired-end short read Hi-C data ^61^, we used the Hi-C phasing mode of hifiasm (GM19434, HG02282, HG02769, HG02953, HG03452, HG03520). For the remaining 4 samples, hifiasm was run without additional data types (HG00171, HG00864, NA19036, NA20847). Six of the samples (GM19129, HG02282, HG02769, HG02953, HG03452, HG03520) have not been reported on by the HGSVC previously. The average CPU time of hifiasm was 323h at a peak memory usage of 105Gb.

Additionally, four hifiasm-based great-ape assemblies (Bonobo, Chimpanzee, Gorilla and Orangutan) were acquired from a recent publication ^22^. (available at https://doi.org/10.5281/zenodo.4721957). In these four datasets, contigs labeled as ‘primary’ were used.

### Core-duplicons

sSV loci were tested for overlap with genes and gene families mapping to ‘core-duplicons’ ^47^. To create a table of relevant genes, we assembled a list of 22 prominent core duplicon - associated gene families (*NBPF, RANBP2, RGPD, PMS2, PPY, C9orf36, ZNF790, SPRYD5, NPIP* (also known as *Morpheus*), *GOLGA, LRRC37, TBC1D3, USP6, SMN, CCDC127, TRIM51, GUSBP, FAM75A, SPATA31, OR7E, DPY19, SPYDE*). ^62,63^. We subsequently queried the gencode gene annotation (version 35) ^64^ for all members of those families, leading to a list of 131 gene instances (Table S3).

### Automated sequence validation with Nanopore reads

We used ONT reads generated by the HGSV Consortium for 11/26 samples - GM19129, HG00512, HG00731, HG00733, HG02282, HG02769, HG02818, HG02953, HG03452, HG03520, NA19239 (data availability). After removing adapters with Porechop (https://github.com/rrwick/Porechop), ONT reads were mapped to both haplotypes of their respective samples (i.e., reads from each sample were mapped both to the h1 and h2 assembly). We invoked the sniffles2 ^36^ SV caller to identify homozygous variants which correspond to assembly regions not supported by any reads, i.e. likely assembly errors. Using the liftover coordinates provided by NAHRwhals, we queried the sequences corresponding to their CHM13-T2T-counterpart individually for each assembly.

### Validation of four morbid CNV-associated regions with Nanopore reads

Four disease-associated regions were additionally verified using a manual approach. For this, aligned ONT reads were phased post-hoc using ‘samtools phase’ and visualized in IGV, highlighting discordant reads which can be indicative of large-scale rearrangements (Fig. S19).

## Data availability

PacBio HiFi sequencing data, Strand-seq as well as Oxford Nanopore sequencing data were generated by the HGSVC consortium and can be accessed through the HGSVC data portal https://www.internationalgenome.org/data-portal/data-collection/structural-variation. Assembled genomes can be accessed via https://doi.org/10.5281/zenodo.7635935.

## Acknowledgements

We thank Celia Tsapalou, Thomas Weber and Michael Jendrusch for valuable feedback on the code and the manuscript, and the members behind the deutsch_studio account on fiverr.com for designing our logo.

## Contributions

Conceptualization and Methodology were envisioned by W.H., J.O.K and F.S. Development of the Software and subsequent formal analysis was performed by W.H with supervision by F.S and J.O.K. Phased human genome assemblies from long-read genomes generated by the Human Genome Structural Variation Consortium were created by P.E. and T.R. Writing was performed jointly by W.H, J.O.K and F.S with input from all authors.

## Consortia

The members of the Human Genome Structural Variation Consortium (HGSVC) are (*denotes co-chairs): Haley J. Abel, Hufsah Ashraf, Peter A. Audano, Anna O. Basile, Christine Beck, Marc Jan Bonder, Harrison Brand, Marta Byrska-Bishop, Mark J.P. Chaisson, Yu Chen, Ken Chen, Zechen Chong, Nelson T. Chuang, Wayne E. Clarke, André Corvelo, Scott E. Devine, Peter Ebert, Jana Ebler, Evan E. Eichler*, Uday S. Evani, Susan Fairley, Paul Flicek, Sky Gao, Mark B. Gerstein, Maryam Ghareghani, Ira M. Hall, Pille Hallast, William T. Harvey, Patrick Hasenfeld, Alex R. Hastie, Wolfram Höps, PingHsun Hsieh, Sarah Hunt, Jan O. Korbel*, Sushant Kumar, Charles Lee*, Alexandra P. Lewis, Chong Li, Bin Li, Yang I. Li, Jiadong Lin, Tsung-Yu Lu, Rebecca Serra Mari, Tobias Marschall*, Ryan E. Mills, Zepeng Mu, Katherine M. Munson, David Porubsky, Benjamin Raeder, Tobias Rausch, Allison A. Regier, Jingwen Ren, Bernardo Rodriguez-Martin, Ashley D. Sanders, Martin Santamarina, Xinghua Shi, Chen Song, Oliver Stegle, Michael E. Talkowski, Luke J. Tallon, Jose M.C. Tubio, Aaron M. Wenger, Xiaofei Yang, Kai Ye, Feyza Yilmaz, Xuefang Zhao, Weichen Zhou, Qihui Zhu, and Michael C. Zody.

