## Supplemental Figures for "Impact and characterization of serial structural variations across humans and great apes"

### Supplementary Figures

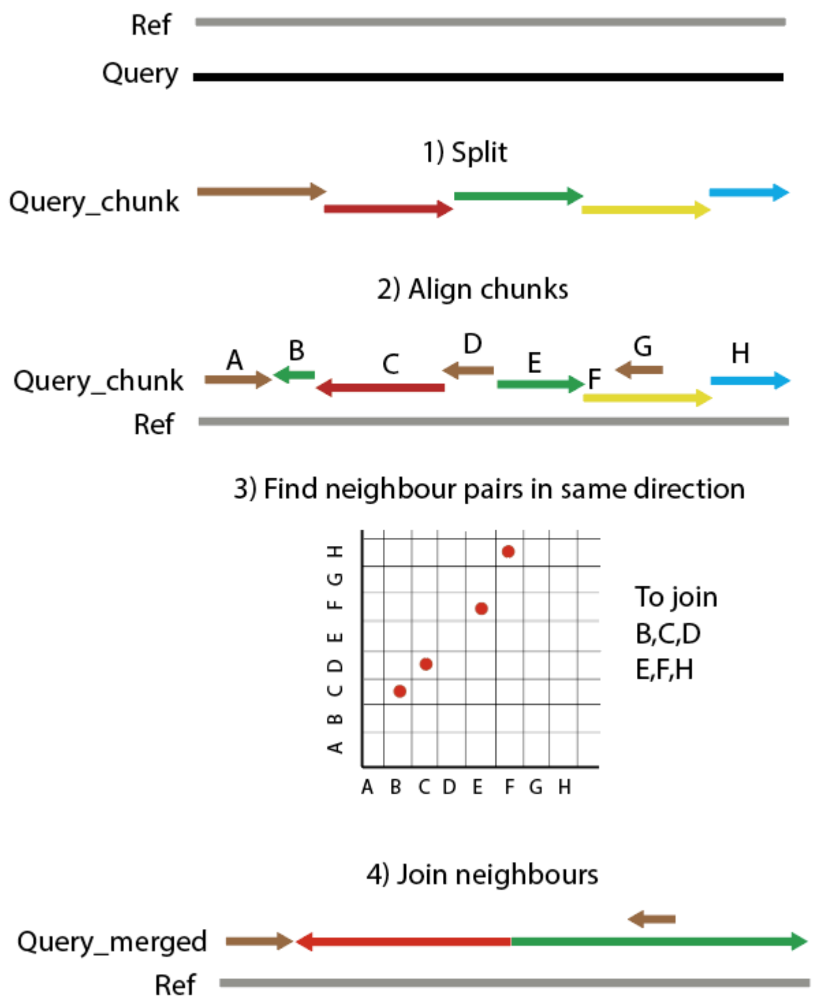


**Figure S1. Schematic view of the custom alignment pipeline. (1)** Before alignment, the Query sequence is split up into equality sized sequence 'chunks' of predefined length (100bp for sequences below 10 kbp, 1 kbp for sequences below 100 kbp, 10 kbp for sequences below 5 Mbp and 20 kbp above). **(2)** The chunks are aligned to the reference individually, using minimap2 aligner (version 2.18-r1035) (Li 2018) (methods). **(3)** Start- and end positions of all-vs-all alignments are then compared to identify pairs where the end of one coincides with the start of another alignment (methods). **(4)** all alignment pairs are then iteratively concatenated to create the final pairwise alignment.

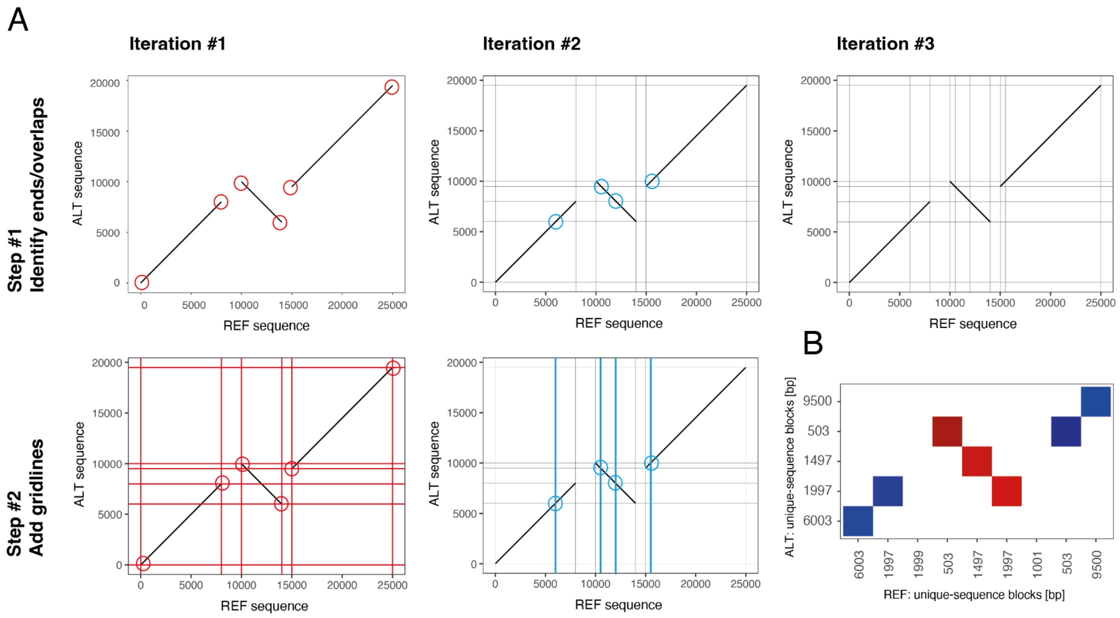


**Figure S2. Visual representation of the iterative dotplot segmentation algorithm. A** Starting from a pairwise alignment, initially all start-and endpoints are identified, and the x- and y-values are noted as the first set of 'gridlines' separating unique alignments. In each subsequent step, novel overlaps between existing gridlines and the pairwise alignments are identified, and subsequently new gridlines are inserted in x and y direction at the intersections. Once the grid has converged, each field is, by design, traversed by zero or one alignment vectors diagonally, intersecting with exactly two opposite corners. Grids which do not converge after 10 iterations are rejected and the dotplot pre-processing is repeated with another parameter set until a converging representation is found (Methods). **B** Using the determined grid as a reference, we derive a "condensed dotplot" where each field of the new dotplot represents a sector of the grid, the value represents the length of the traversing alignment, and the sign of the value corresponds to the direction of the alignment (blue: positive values: direct orientation; red: negative values: inverse orientation).

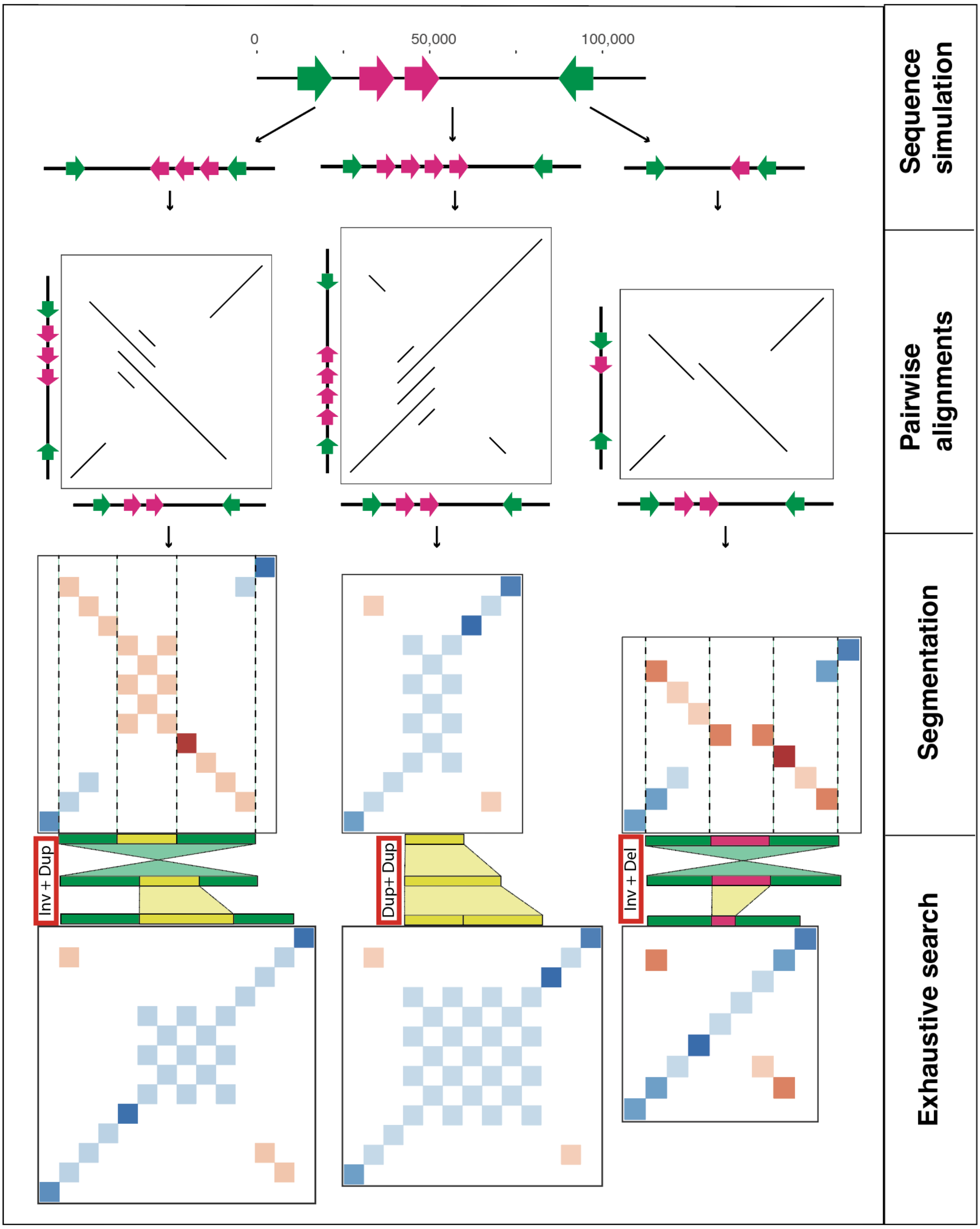


**Figure S3. Workflow of simulation experiments.** For each iteration in the simulation experiments, an artificial genomic region was created with two pairs of SDs, length and similarity of SDs as pre-defined parameters, and the position and orientation of SDs was randomly chosen each time. Simulated sequences were then subjected to in-silico mutation, applying all possible NAHR-chains of depth 1 and 2, resulting in three to >10 mutated ‘alternative’ alleles derived from one founder sequence. Each alternative sequence was tested for mutations using the NAHRwhal algorithm consisting of pairwise alignments, dotplot segmentation and exhaustive search. 50 genomic founder sequences were created for each combination of three SD similarities (90%, 95%, 99%) and four different SD lengths (100 bp, 500 bp, 1.000 bp, 10.000 bp). The resulting SV calls were finally compared to the in-silico mutation ground truth.

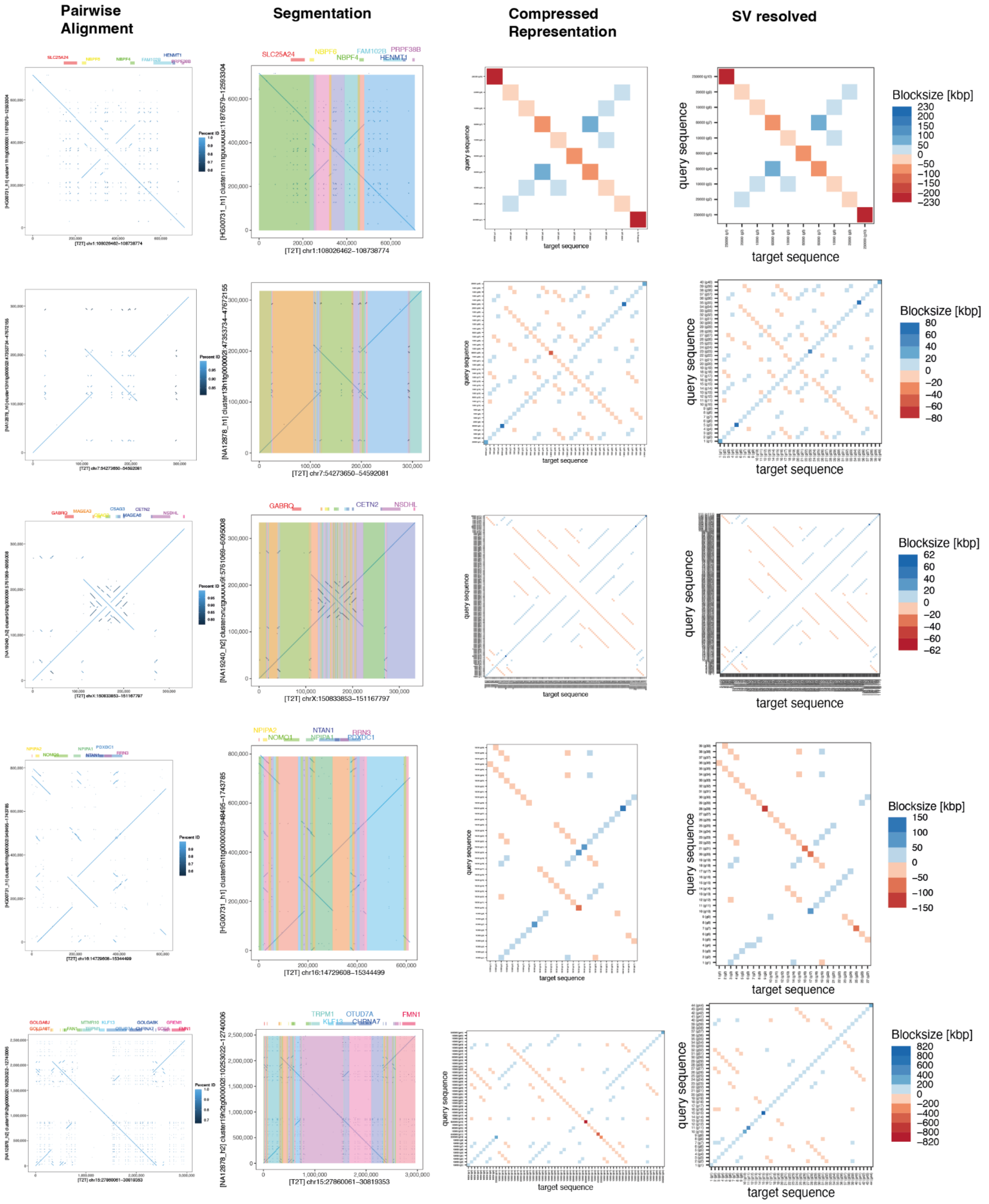


**Figure S4. Unittest loci 1-5/10.** Pairwise alignments (left), segmentations (plotted on x-axis only for clarity), condensed dotplots and mutation-resolved condensed dotplots for five example loci representing loci of different SD- and SV complexity. The segmentation algorithm scales window size dynamically depending on the complexity of a locus, leading to a fine representation of complex, repetitive regions in the compressed representation.

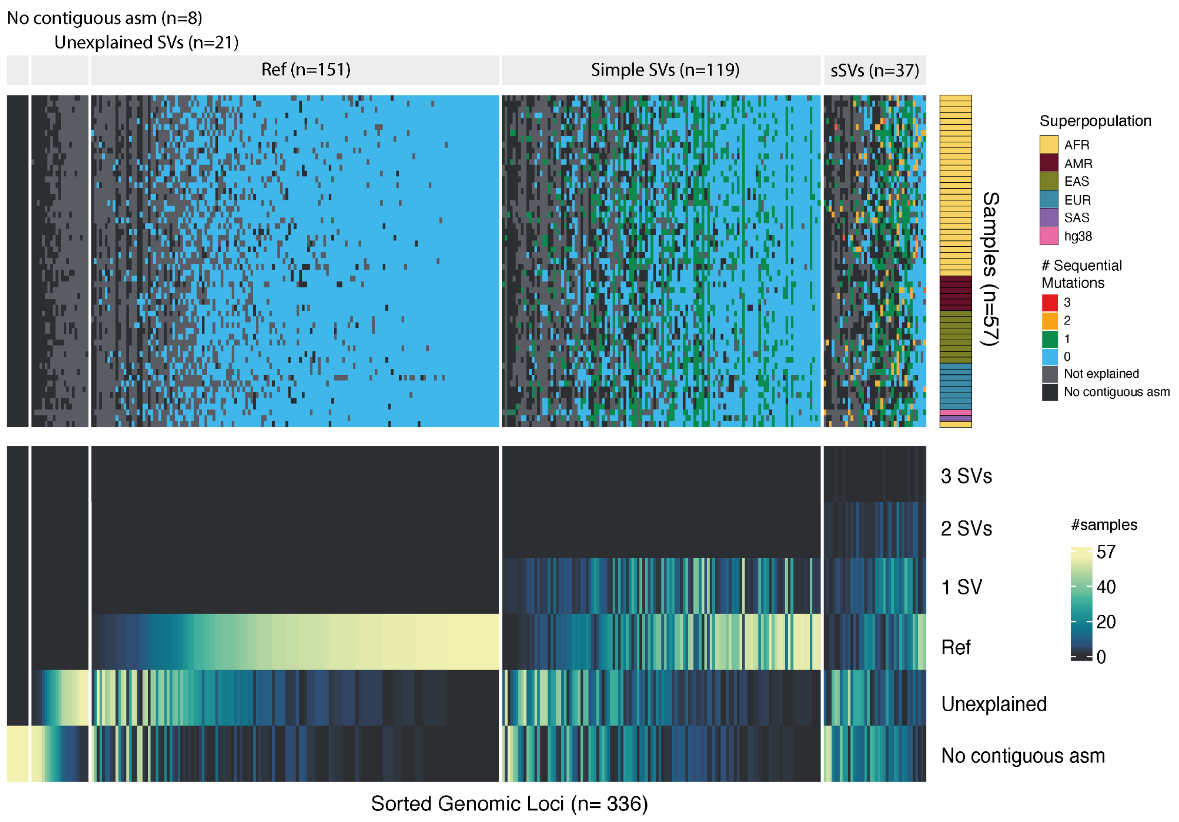


**Figure S5. Detailed results of scanning 336 loci across 57 haplotypes with NAHRwhals. (Top)** Visualization of every SV call per sample and locus. Loci were grouped according to the number of samples displaying No contiguous assembly, Unexplained SVs and mutations of various depth. Sample ancestry is indicated on the right. **(Bottom)** Simplified view representing the number of various results per locus. n=37 loci displayed nested SVs ('2 SVs', '3SVs') in at least one sample. In cases where loci contained >1 non-overlapping simple SVs, these were reported as '1 SV', reflective of their maximum depth.

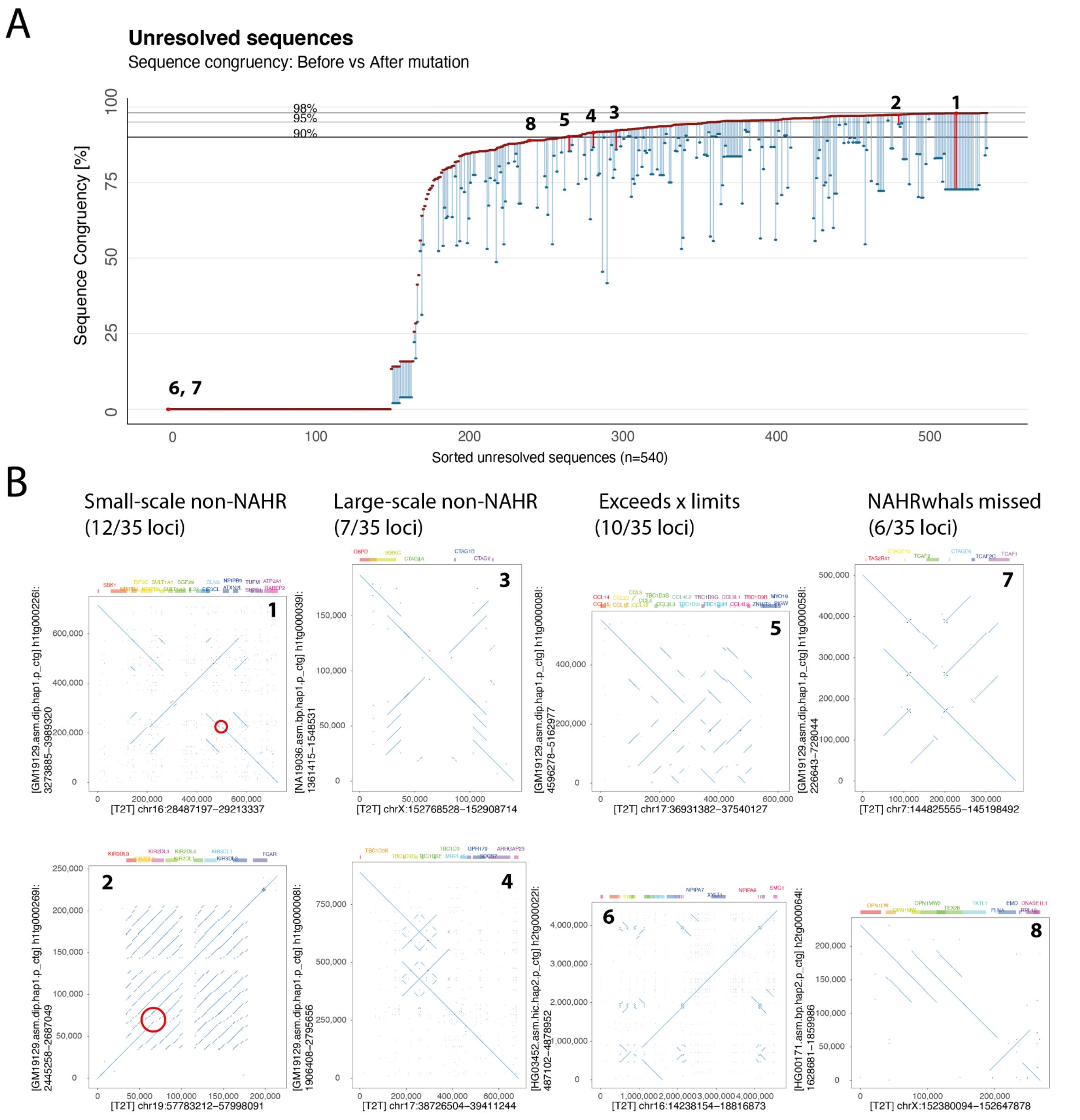


**Figure S6.** Investigation of variant loci flagged as 'unexplained' by NAHRwhals**. A** A dumbbell plot showing the sequence congruency between each of the 540 unexplained sequences and the CHM13-T2T reference before (blue dots) and after (red dots) application of the highest-scoring mutation chain determined by NAHRwhals. Numerals indicate alignments followed up in panel B. **B** Examples of unexplained alignments determined by manual investigation of a random subset of sequences depicted in A. Unexplained loci group into small-scale non-NAHR events, large-scale non-NAHR events, Mutations exceeding our chosen reference locus and NAHR-mutations which were missed by NAHRwhals.

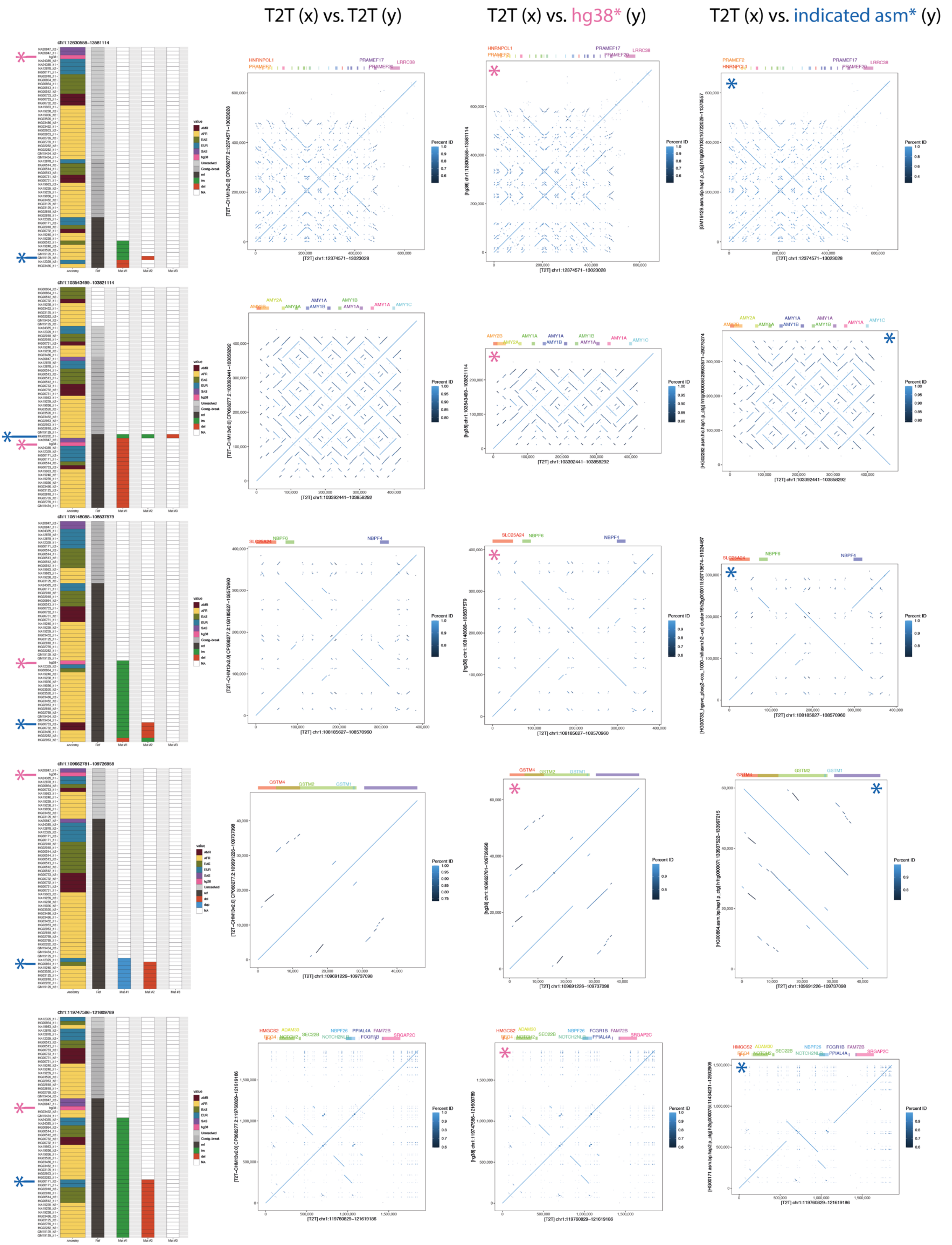
T2T (x) vs. T2T (y)
T2T (x) vs. hg38* (y)
T2T (x) vs. indicated asm* (y)
chr1:12830558–13581114
chr1:103543499–103821114
chr1:108148088–108537579
chr1:109662781–109726958
chr1:119747586–121609789
HNRNPCL1
PRAMEF2
PRAMEF17
PRAMEF20
LRRC38
AMY2A
AMY2B
AMY1A
AMY1B
AMY1C
SLC25A24
NBPF6
NBPF4
GSTM4
GSTM2
GSTM1
HMGCS2
ADAM30
SEC22B
NBPF26
PPIAL4A
FAM72B
SRGAP2C
FCGR1B
NOTCH2NL
Percent ID
[T2T] chr1:12374571–13023028
[T2T] chr1:103392441–103858292
[T2T] chr1:108185627–108570960
[T2T] chr1:109691226–109737098
[T2T] chr1:119760829–121619186
Ancestry
Ref
Mat #1
Mat #2
Mat #3

**Figure S7. sSV calls and dotplot visualizations of the first 5/37 human sSV loci by start coordinate.** For each locus, we display 1) the full SV callset (left), 2) the CHM13-T2T reference aligned to itself, 3) the CHM13-T2T reference aligned to hg38, 4) the CHM13-T2T reference aligned to the human assembly with the longest called mutational chain. If multiple chains have similar length, we default to displaying the sample with the highest y-value in the sSV-calls plot (left). Colored asterisks are overlaid for visual guidance between genotypes and dot plots. Assembly contigs are, by chance, denoted as the reverse complement to the CHM13-T2T reference in approx. 50% of cases (here: loci 1-4). The directionality of the y axis in the dotplots is therefore arbitrary.

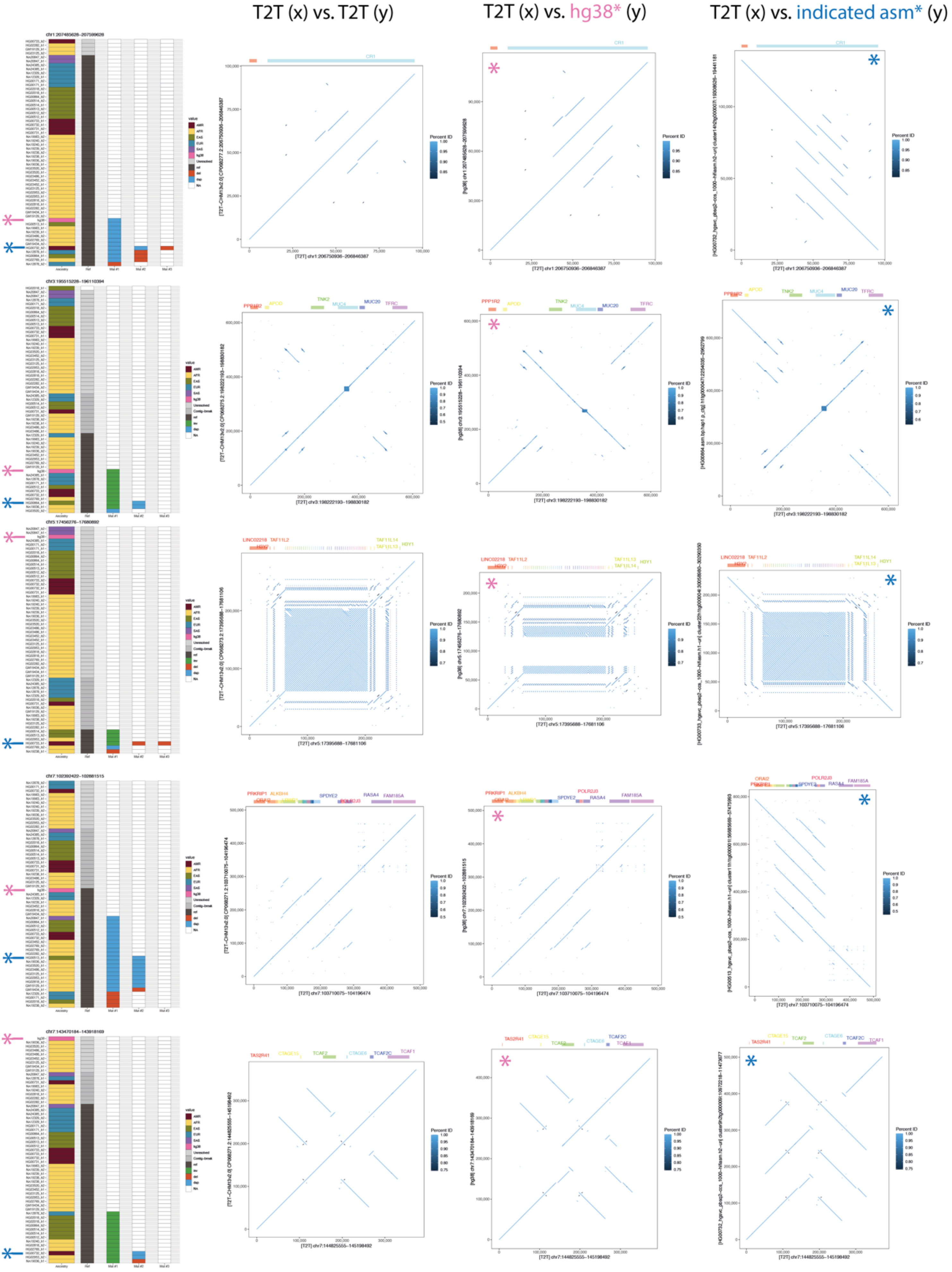


**Figure S8.** Continuation of **Figure S7.**

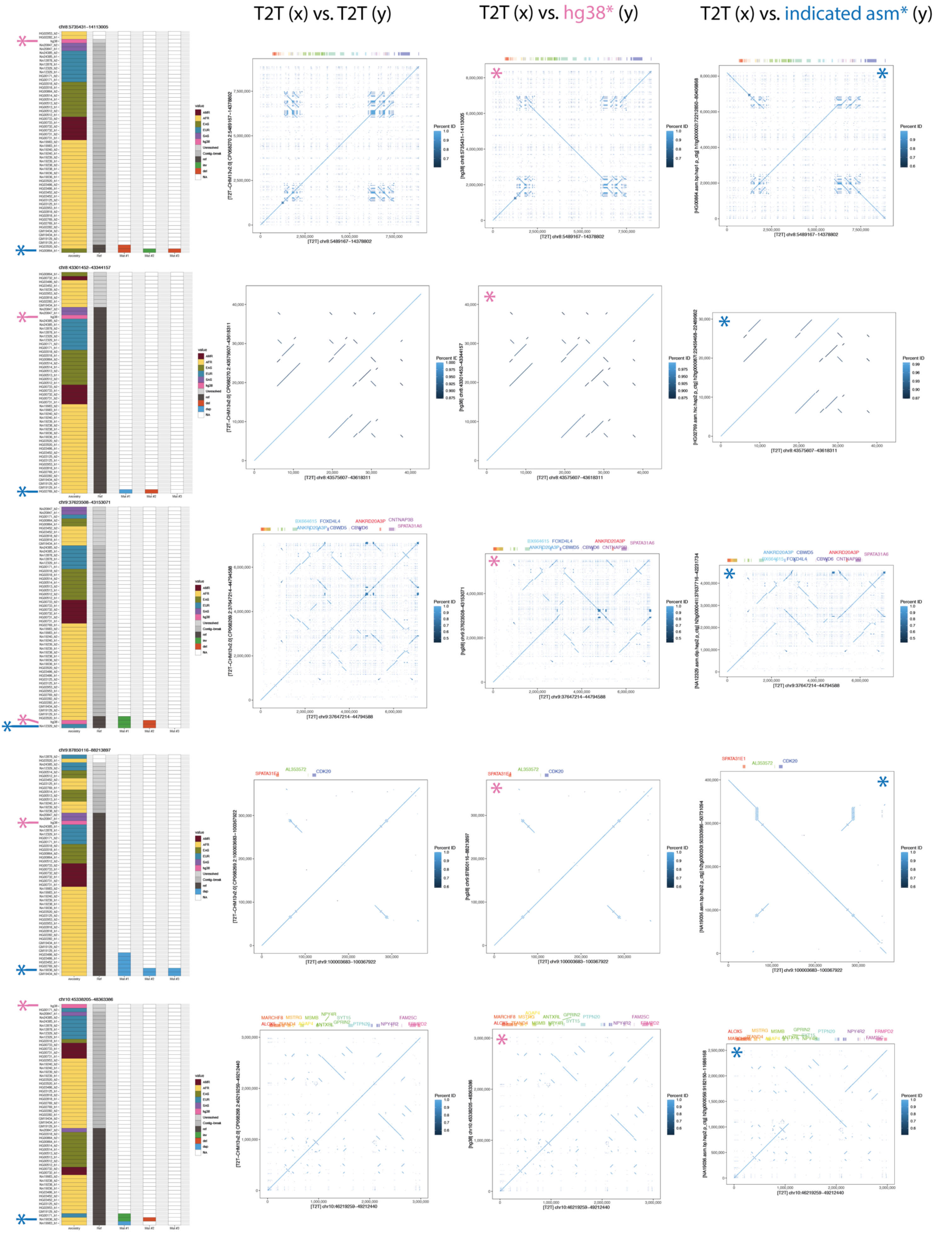


**Figure S9.** Continuation of **Figure S8.**

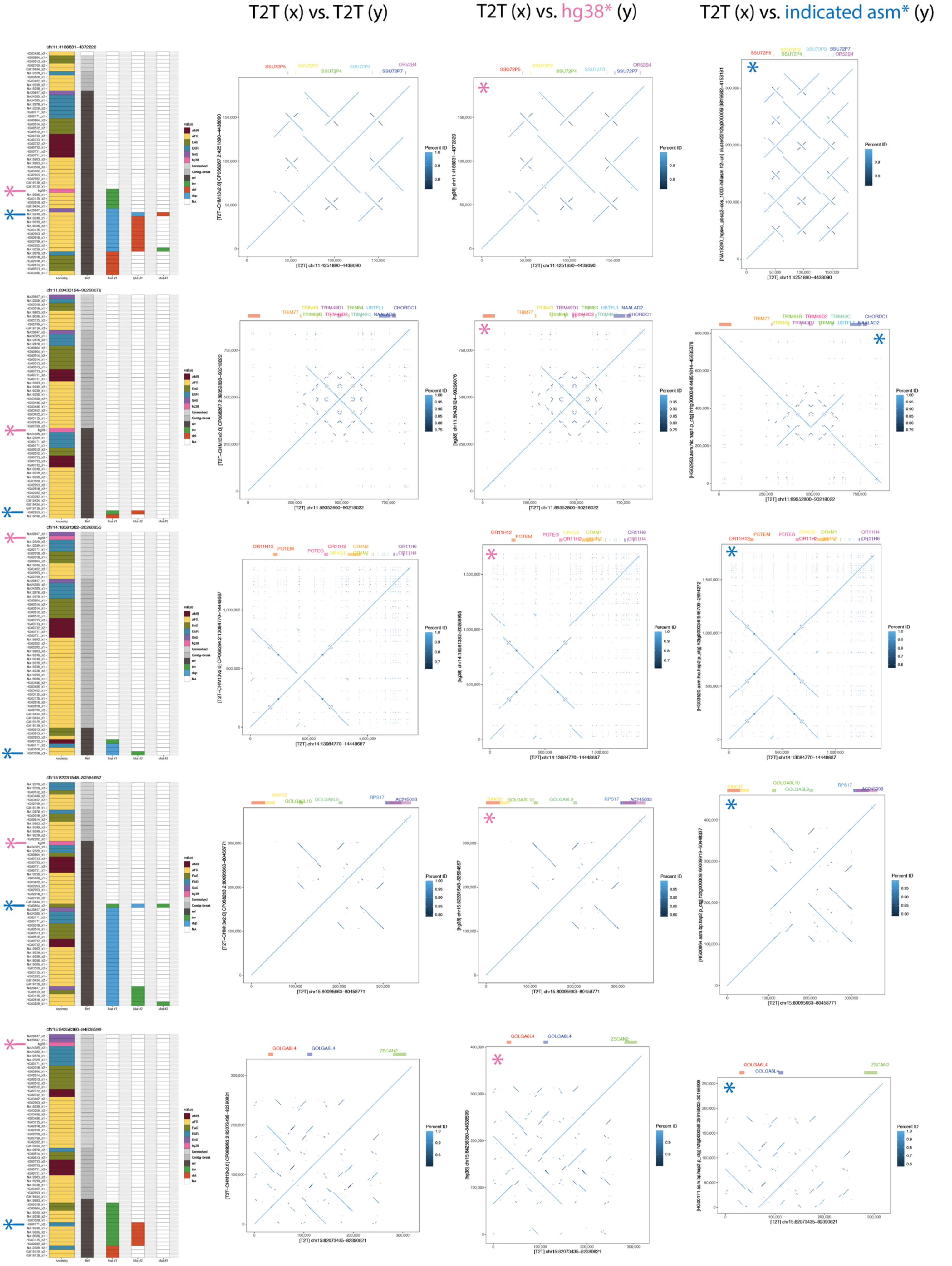


**Figure S10.** Continuation of **Figure S9.**

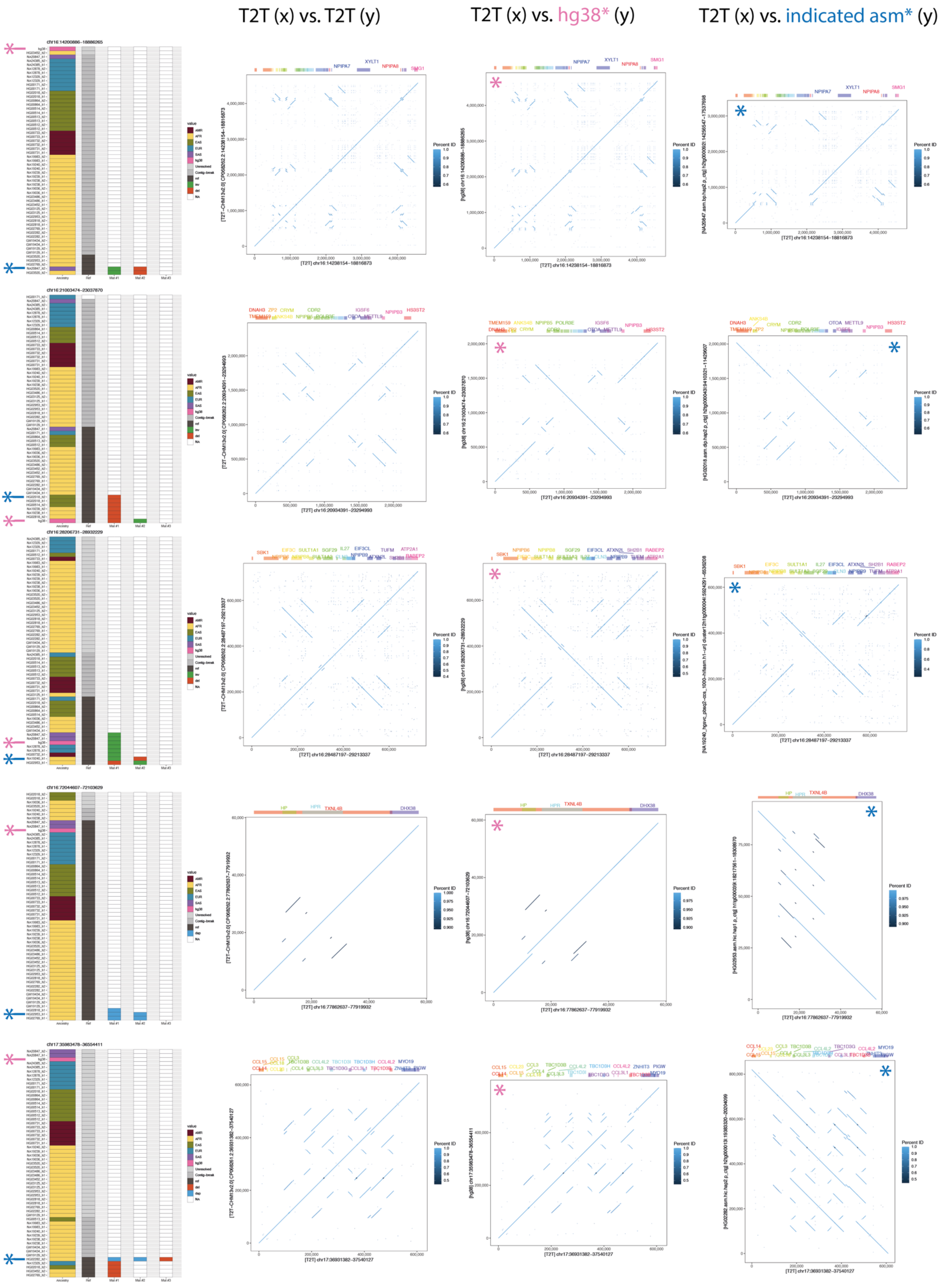


**Figure S11.** Continuation of **Figure S10.**

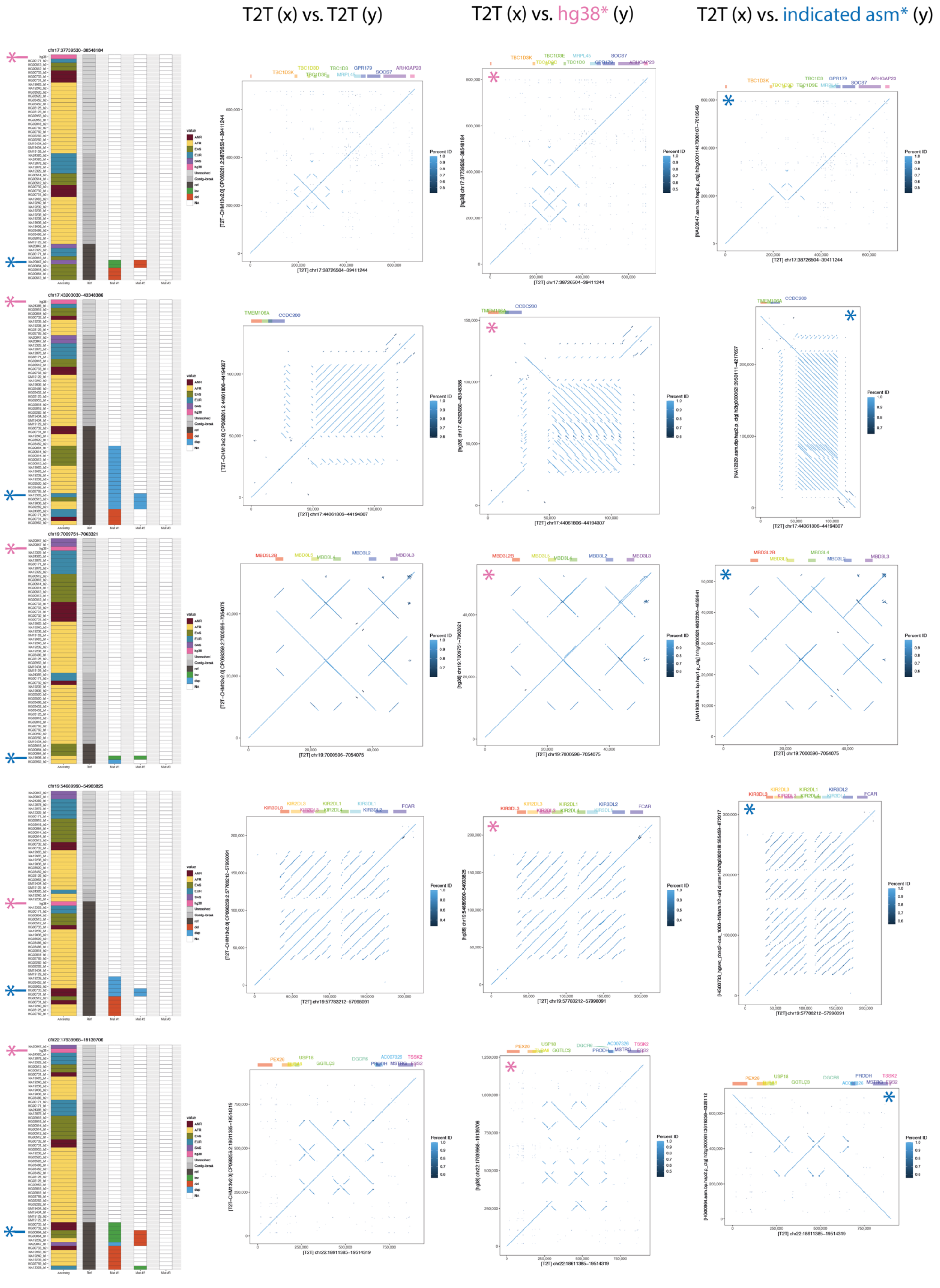


**Figure S12.** Continuation of **Figure S11.**

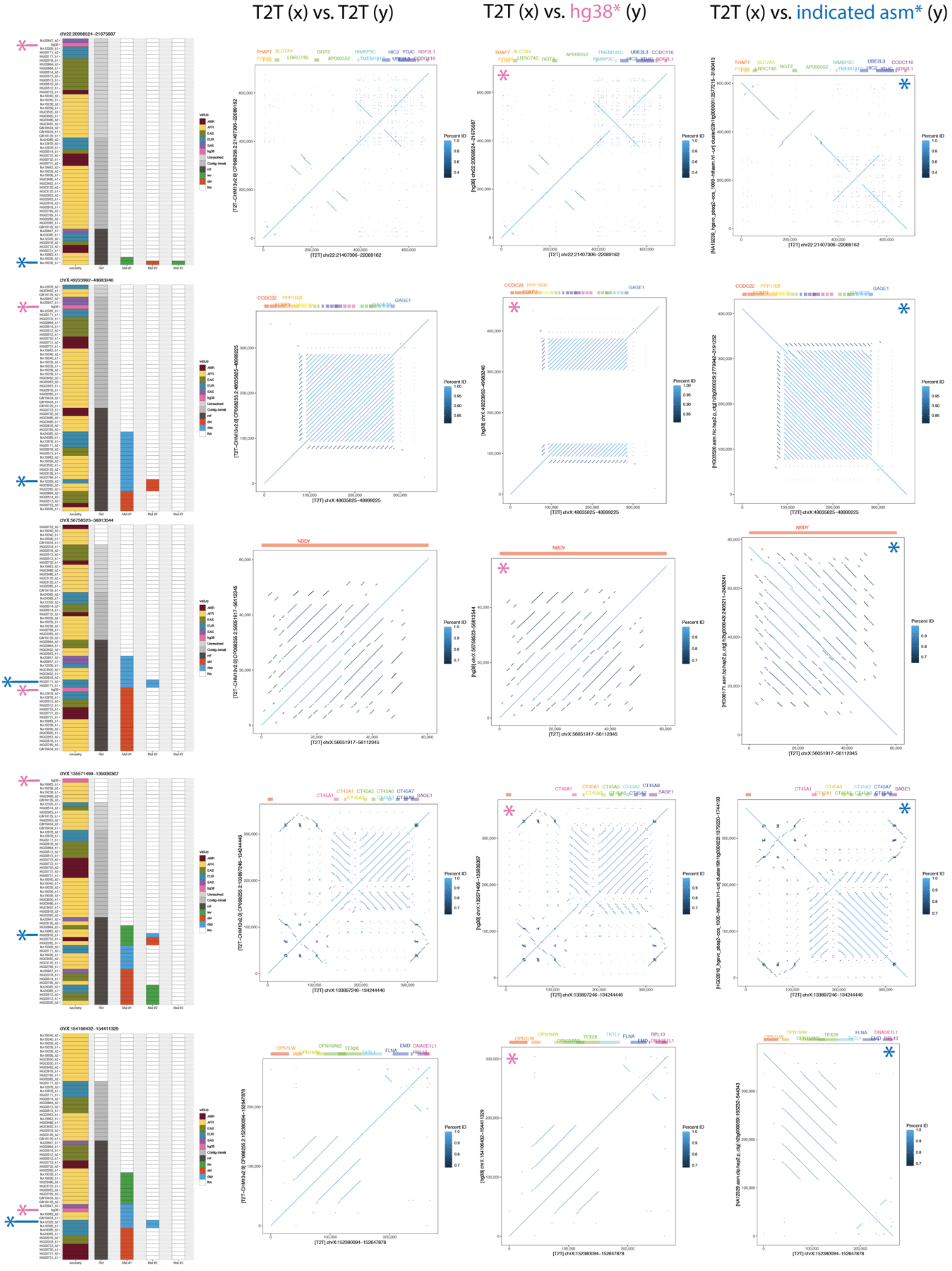


**Figure S13.** Continuation of **Figure S12.**

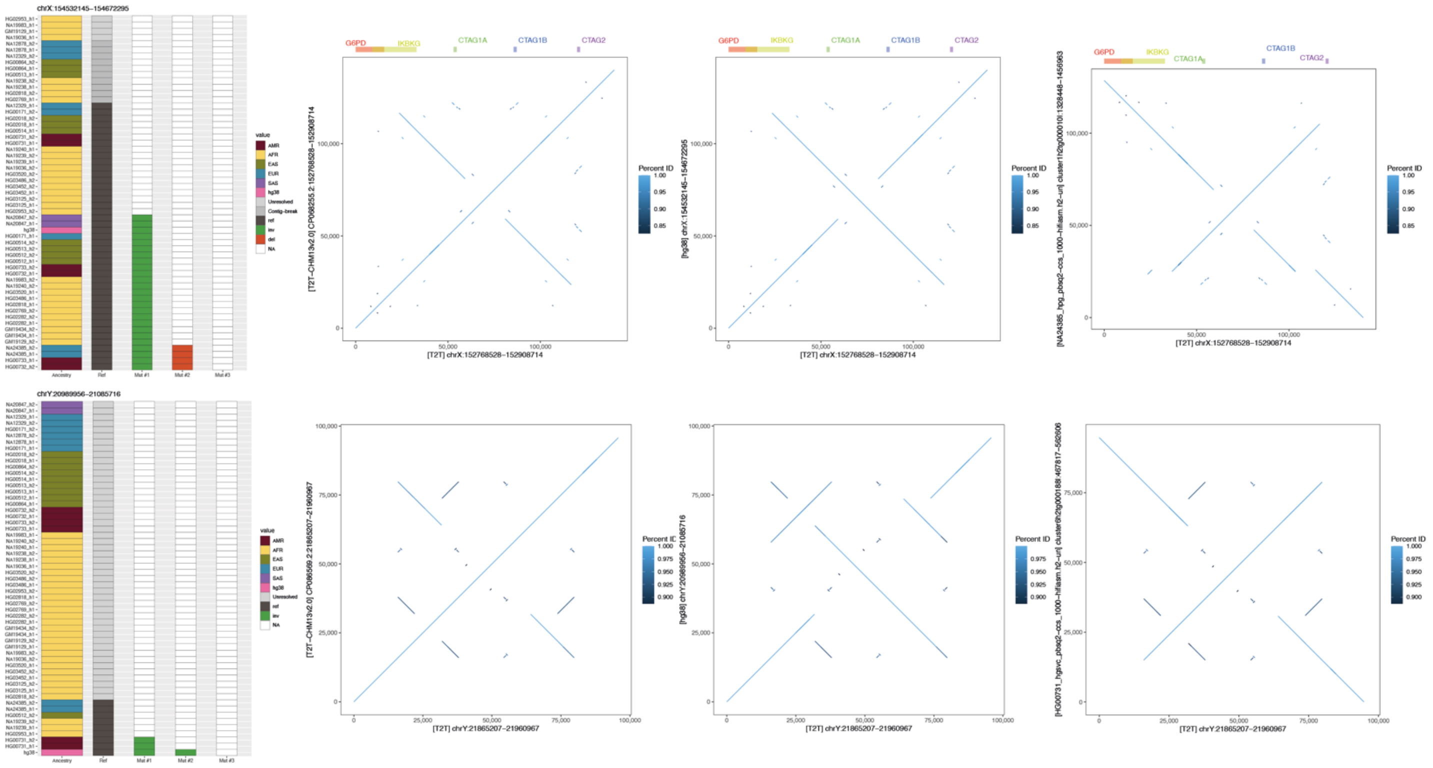

**Figure S14.** Continuation of **Figure S13.**

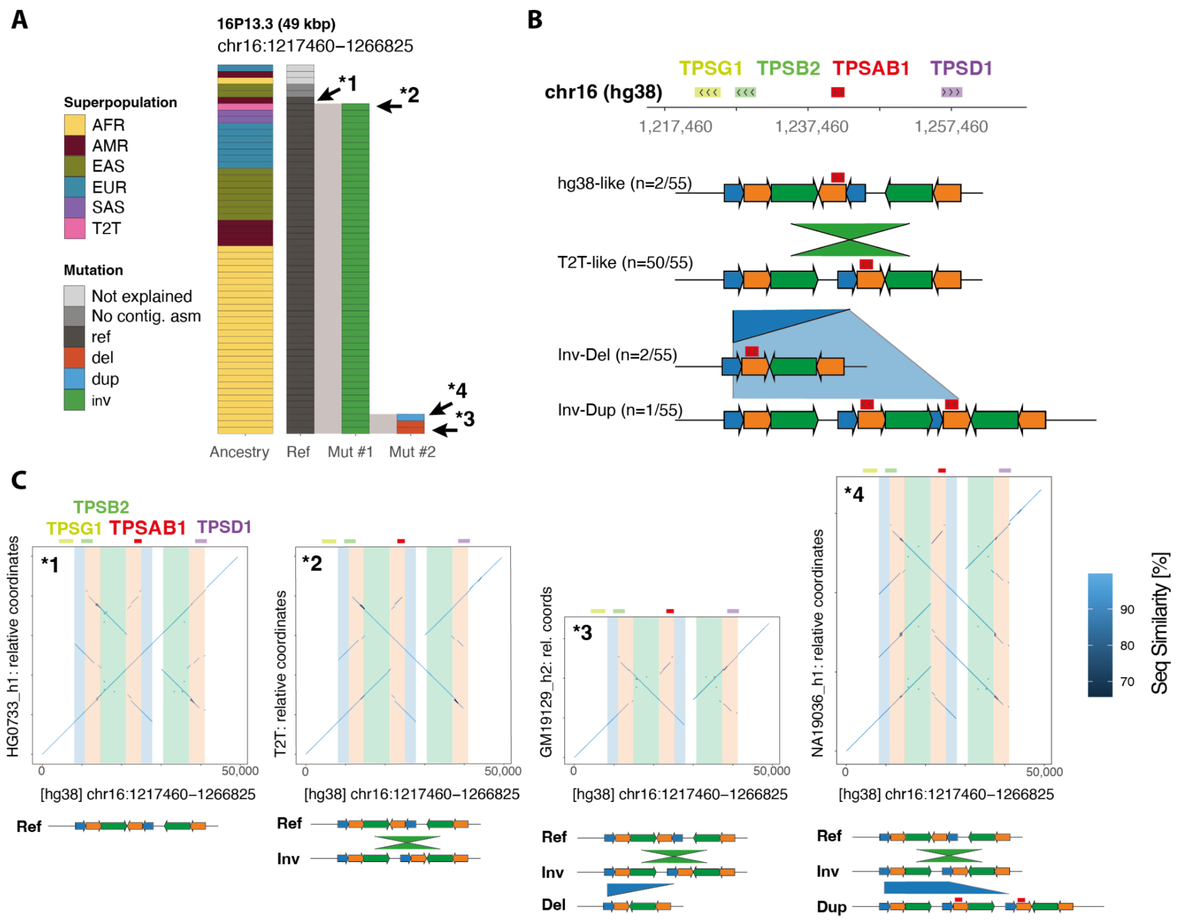


**Figure S15. Hg38-exclusive sSVs in the TPSAB1-containing locus. A** Mutations identified in the TPSAB1-containing region on chr16p13.3 with respect to the hg38 reference. The majority of haplotypes carry the inverted allele. **B** Schematic of the inferred sSV configurations **C** Dotplots visualising four distinct haplotype configurations, two of which carry potentially functional CNVs.

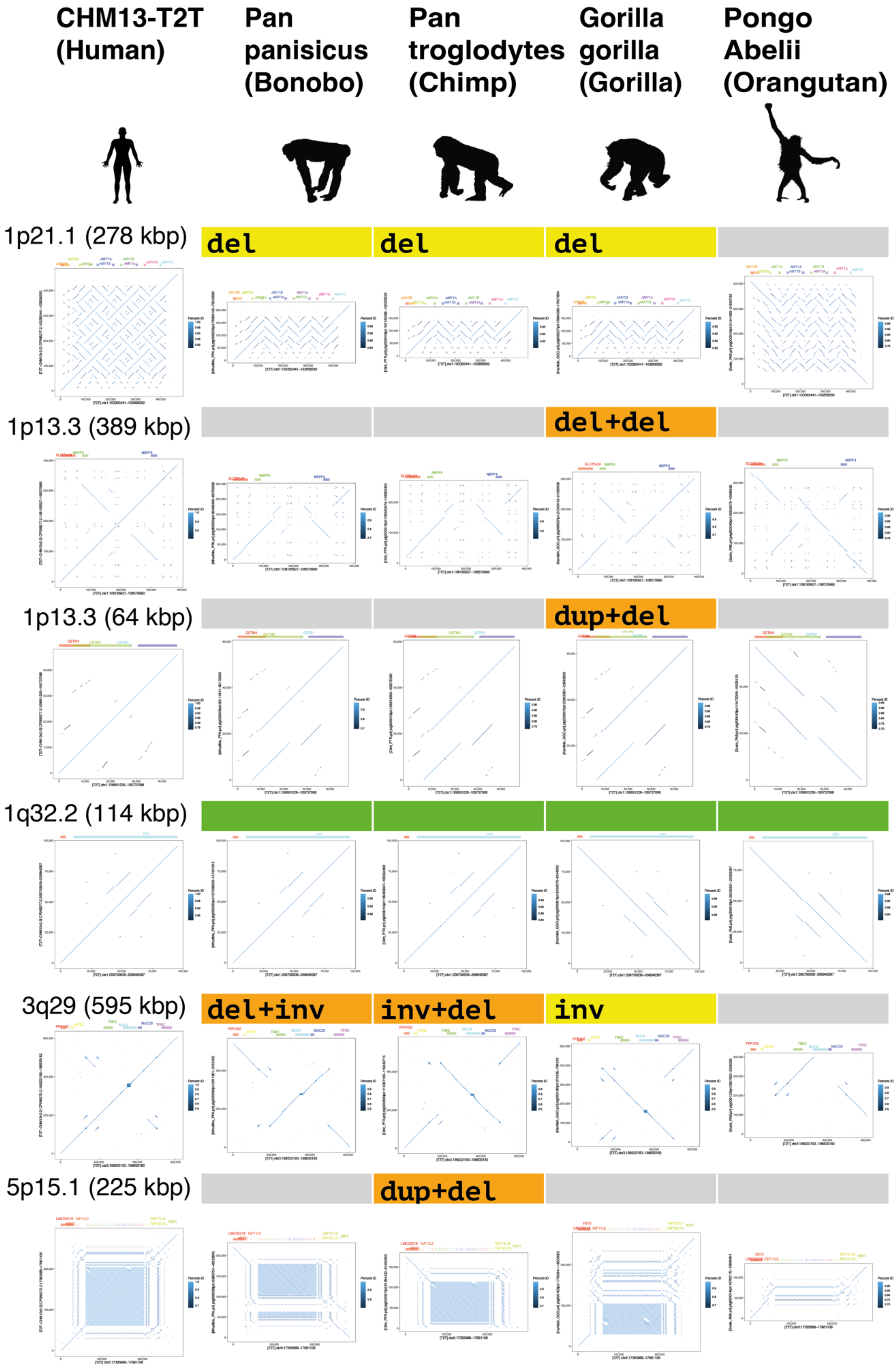
CHM13-T2T (Human)
Pan panisicus (Bonobo)
Pan troglodytes (Chimp)
Gorilla gorilla (Gorilla)
Pongo Abelii (Orangutan)
1p21.1 (278 kbp)
del
del
del
1p13.3 (389 kbp)
del+del
1p13.3 (64 kbp)
dup+del
1q32.2 (114 kbp)
3q29 (595 kbp)
del+inv
inv+del
inv
5p15.1 (225 kbp)
dup+del

**Figure S16. Dotplots of 16 human sSV regions reported as ‘resolved’ in at least one great ape assembly.** Reported genotypes (relative to CHM13-T2T) are indicated above each dotplot. The directionality of the y axis in the dotplots is arbitrary given the random orientation (direct vs reverse complement) of contigs. Green: reference. Light grey: unexplained variance.

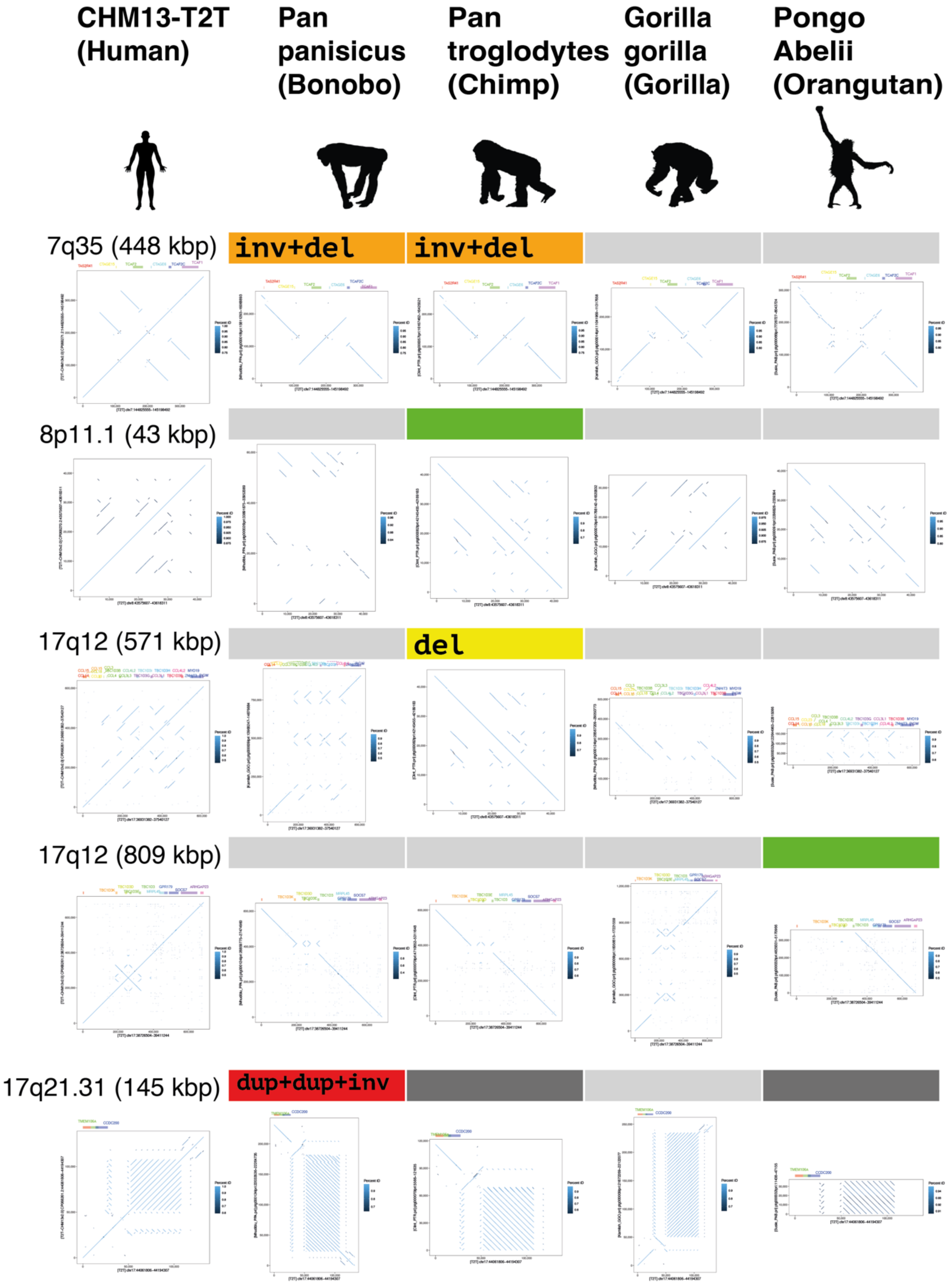


**Figure S17. Continuation of Figure S16.** Dark grey: non-continuous assembly.

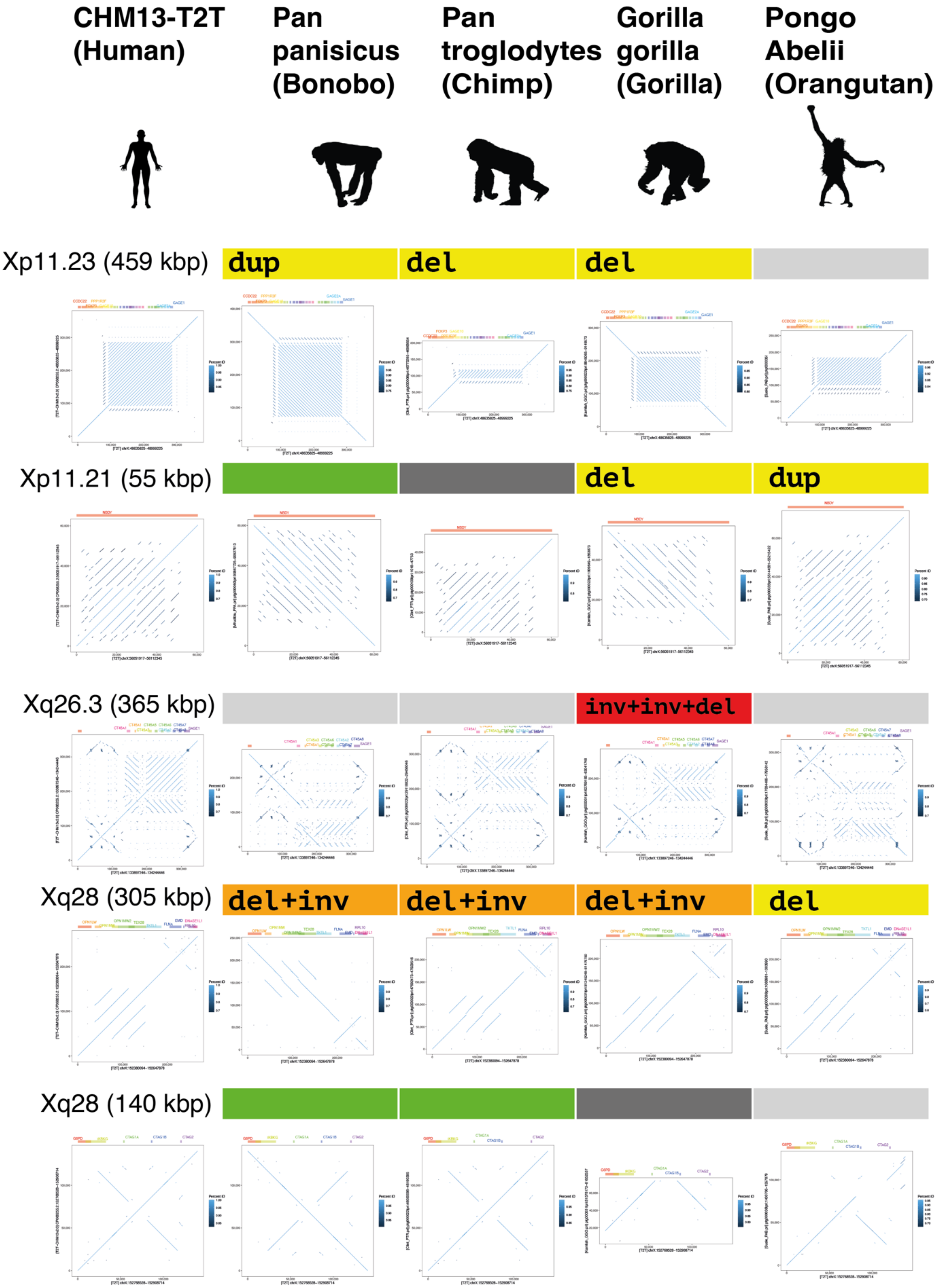


**Figure S18. Continuation of Figure S17.**

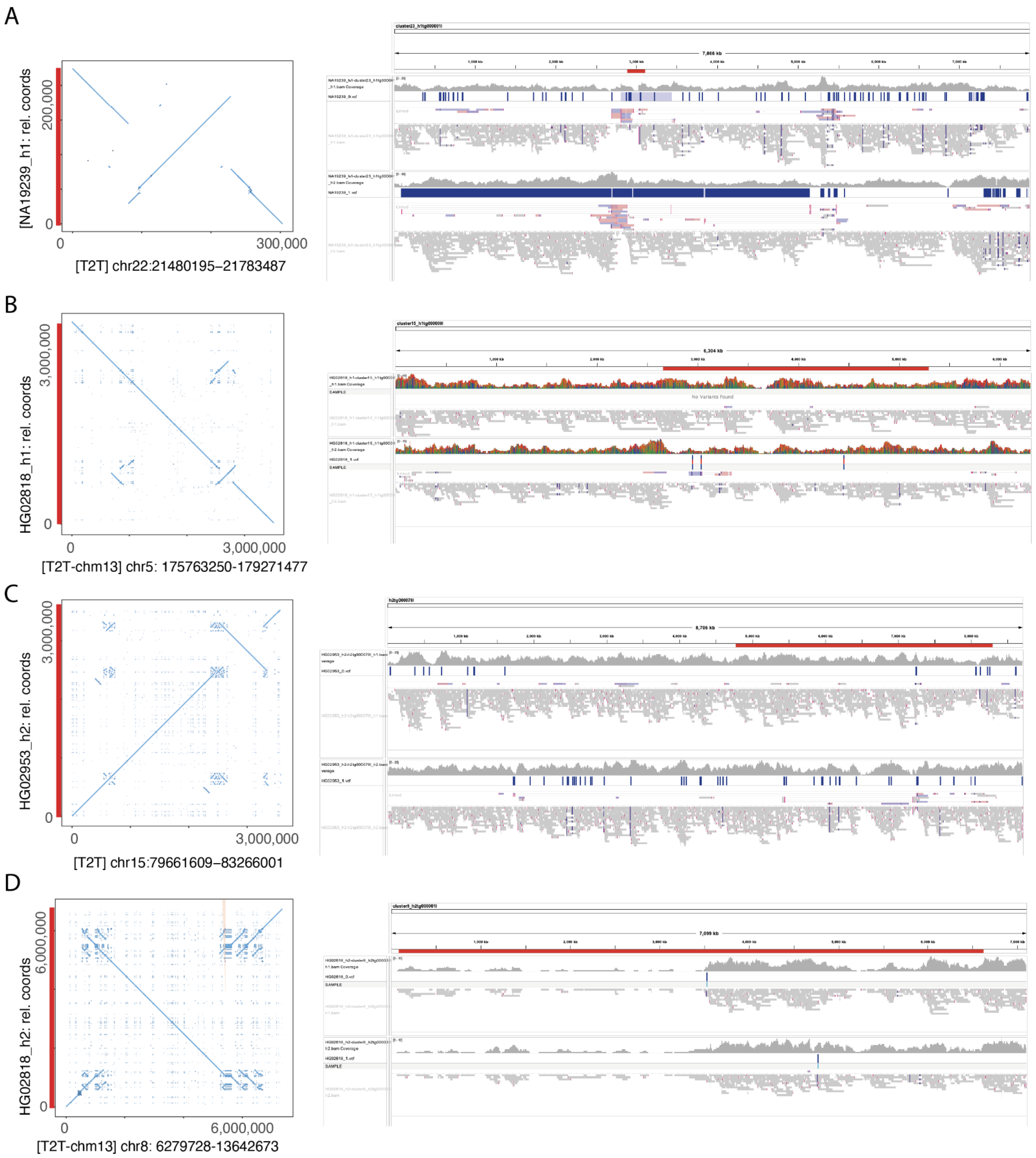


**Figure S19. ONT reads aligned to their respective assemblies in four morbid CNV - containing regions.** IGV screenshots of nanopore reads mapping to the entire assembled contig are depicted on the right, with a red highlight bar indicating the position of the window region of interest. Sniffles-based calls per haplotype are overlaid over each read track. Reads were split into haplotypes h1 (top track per panel) and h2 (bottom track per panel) using samtools phase. The depicted regions are **A** Di-George syndrome region, **B** Sotos syndrome region, **C** 15q25.2 del/dup region, **D** 8q23.1 del/dup region.

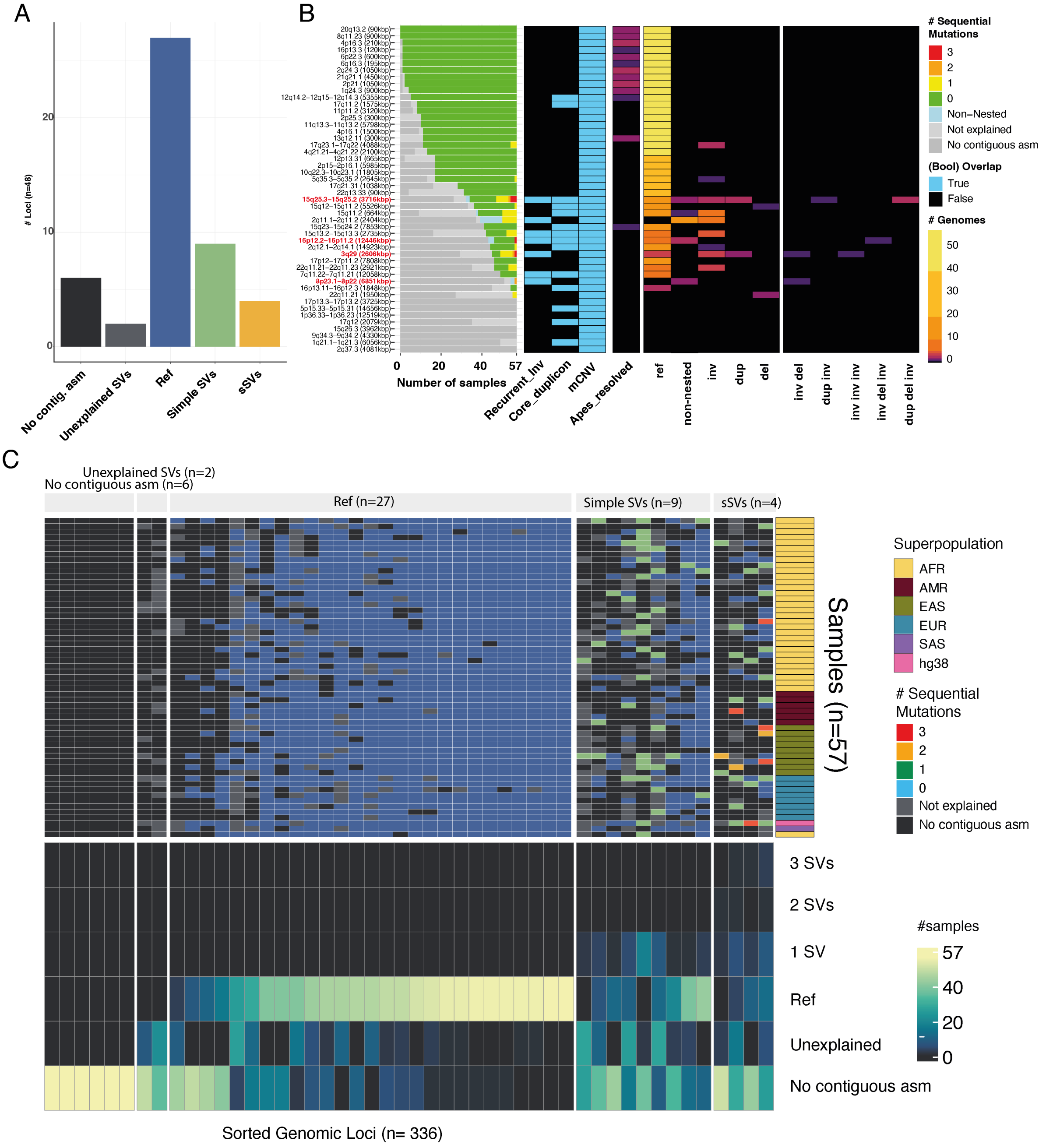


**Figure S20. Overview over the NAHRwhals results of scanning 48 mCNV-associated regions for sSV content. A** Broad classification of the 48 loci surveyed with NAHRwhals. Loci were considered as 'sSVs' if they displayed at least one overlapping pair of SVs in at least one sample. **B** Overview over the full callset of all 48 loci. The diagram shows the prediction performance in humans and apes ('SVs resolved'), the presence of recurrent inversions, core duplicon-mapping genes and morbid CNV regions in the genomic region, as well as genotypes for each locus. **C (Top)** Visualization of every SV call per sample and locus. Loci were grouped according to the number of samples displaying No contiguous assembly, Unexplained SVs and mutations of various depth. Sample ancestry is indicated on the right. **(Bottom)** Simplified view representing the number of various results per locus. n=37 loci displayed nested SVs ('2 SVs', '3SVs') in at least one sample. In cases where loci contained >1 non-overlapping simple SVs, these were reported as '1 SV', reflective of their maximum depth.

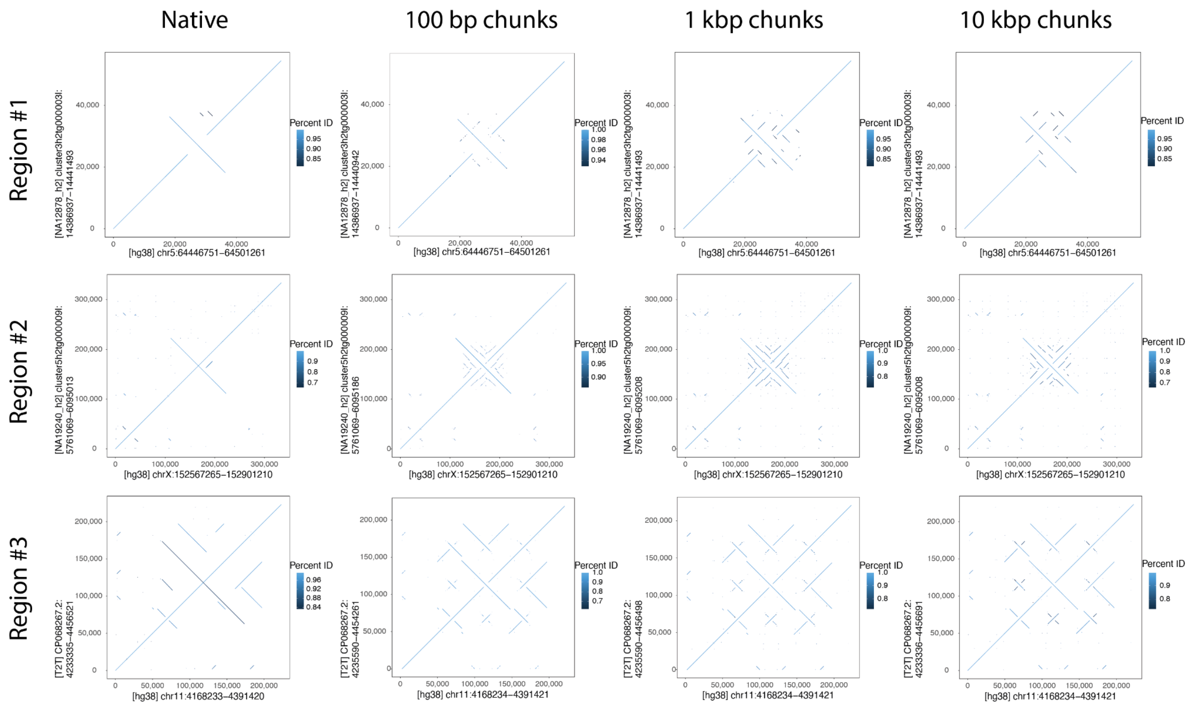


**Figure S21. Three example loci illustrating the effect of chunking input query sequences before alignment.** All panels were created with minimap2 (version 2.18-r1035-dirty; parameters *-x asm20 -P -c -s 0 -M 0.2)*. Without chunking ('Native'), alignments did not resolve all segmental duplications, and reported several inversion regions as palindromes (Regions #2 and #3). Chunking into 100 bp, 1 kbp or 10 kbp reads greatly improves fidelity of the alignments.
